# Cell jamming transitions shape regulatory protein gradients and prime evolutionary divergence

**DOI:** 10.1101/2025.03.05.641736

**Authors:** Alexander V. Badyaev, Cody A. Lee, Maxwell J. Gleason, Georgy A. Semenov, Sarah E. Britton, Carmen Sánchez Moreno, Renée A. Duckworth

**Author notes:** **Author for correspondence:** Alexander Badyaev.

## Abstract

A long-standing goal of evolutionary developmental biology is to identify the mechanisms underlying criticality of developmental transitions that allow processes governing individual cells scale up to the organism-level patterning. The viscoelastic properties of embryonic tissues imply collective cell behaviors, leading to the expectation that signaling networks should capitalize on the material properties of tissues, structuring morphogenesis around the spatial and temporal transitions that they induce. Here, we show that this interaction is evident even prior to tissue differentiation and is traceable to behavior of individual cells. In avian beak primordia, we find that fields of mesenchymal cells undergo cycles of local jamming dynamically modulating coordination of cell shape and movement. These cycles progressively alter the spatial reach of regulatory proteins, strongly expanding or restricting their gradients based on tissue mechanical state. Tissue-level gradients of proteins most sensitive to local cell jamming transitions also diverge the most across populations, priming tissue compartmentalization. These findings suggest that the material state transition is an effective interface for integration of stochastic physical processes and genetic regulation and is well placed to underlie criticality of developmental systems allowing local rules governing cell-state transitions scale up to tissue-level patterning. More broadly, our findings reveal how transient material transitions reset developmental trajectories and promote diversification while preserving robustness.

## 1. Introduction

Morphogenesis of multicellular organisms occurs at the level of tissues but is produced by coordination among thousands of individual cells, requiring integration of distinct processes and consistency of rules across vast spatial and temporal scales. Mechanisms at the interface of physical and biological processes are well suited for this integration: genetic and biochemical controls routinely capitalize on predictable material properties of cells and tissues to produce scale expansion, pattern formation, time measurement, and compartmentalization [1–4].

A prime example of this integration is the transient association of mechanistic and regulatory processes in the jamming transition in cell groups during tissue reorganization in development [5–9]. During early embryogenesis, cells in homogeneous fields routinely undergo spontaneous spatial rearrangements caused by accumulation and dissipation of the compressive stress that cells impose on each other. This process leads to the transient appearance of two tissue states: a glassy-like jammed state of slow-moving cells and a fluid-like unjammed state of fast moving cells. Although bidirectional transitions between jammed and unjammed states is a fundamental component of development, wound repair, and cancer [9–13], how regulatory and cellular processes interact during these transitions and their evolutionary consequences are poorly understood.

In developing tissues, anything that affects cell shape, density, growth, competition, migration, adhesion, or differentiation can trigger jamming transitions [7, 14, 15]. Further, in biological systems, unlike their physical counterparts, jamming transitions can be both active and passive: they can be induced by cells themselves through changes in shape, cycle synchronization, or adhesion propensity, or can be imposed on cells by surrounding tissues of different rigidity or by crowding in a confined space [16–18]. In all of these cases, however, a local process affecting just a few cells leads to larger-scale structural reorganization of the tissue without additional developmental controls at the tissue level [1, 14, 18]. These processes are thus well placed to underlie criticality of developmental systems [19]. This is because they provide links among levels and components of disparate organization and nature (e.g., physical vs biological) in such a way that the regulatory system at each level stabilizes a large-scale expansion arising through incremental variation at the preceding stage without causing these transformations directly. The search for such processes is a long-standing goal of evolutionary developmental biology [20, 21].

The multitude of potential causes of jamming transitions precludes its monopolization by any one biological regulator during evolution. Instead, regulatory networks are expected to evolve to capitalize on the scale-transcending and path-resetting effects of these transitions [22–25]. Further, cell mechanical state and the synthesis of regulatory proteins are directly and reciprocally linked [26, 27], both through the effects of jamming transitions on membrane-to-nucleus distances, cytoplasmic and nuclear biochemistry, and accessibility of cell and nuclear membranes (as a function of cell isomorphy and membrane porosity; [4, 28, 29] and through the effects of tissue material properties on morphogen propagation and concentration [28, 30, 31]. Coupled with the scale-amplifying effects of cell jamming itself, this integration should result in mesoscale consequences for developmental compartmentalization that should also be detectable in evolutionary diversifications.

Here we show that this is indeed the case for ontogeny of beaks across recently established house finch populations [32]. We first document the ubiquity of cell jamming transitions in otherwise homogeneous fields of pre-condensation mesenchyme of upper beak primordia. We then show that the expression and spatial reach of proteins central in avian beak development [33–36] are linked to these jamming cycles. We then test whether cell-jamming associated protein expression primes or buffers early tissue compartmentalization and examine the evolutionary consequences of this association for divergence in protein gradients among recently established populations (Fig. 1). Our findings show how molecular signaling linked with mechanical cell states can be coopted for tissue compartmentalization in morphogenesis, and how conserved biological regulation can integrate with stochastic and transient physical processes to shape evolutionary change. The scale-expansion effect of cell jamming transformations also illuminates the developmental origins of the widespread scaling of animal structures, of which avian beaks are a textbook example [37].

**Fig. 1.**
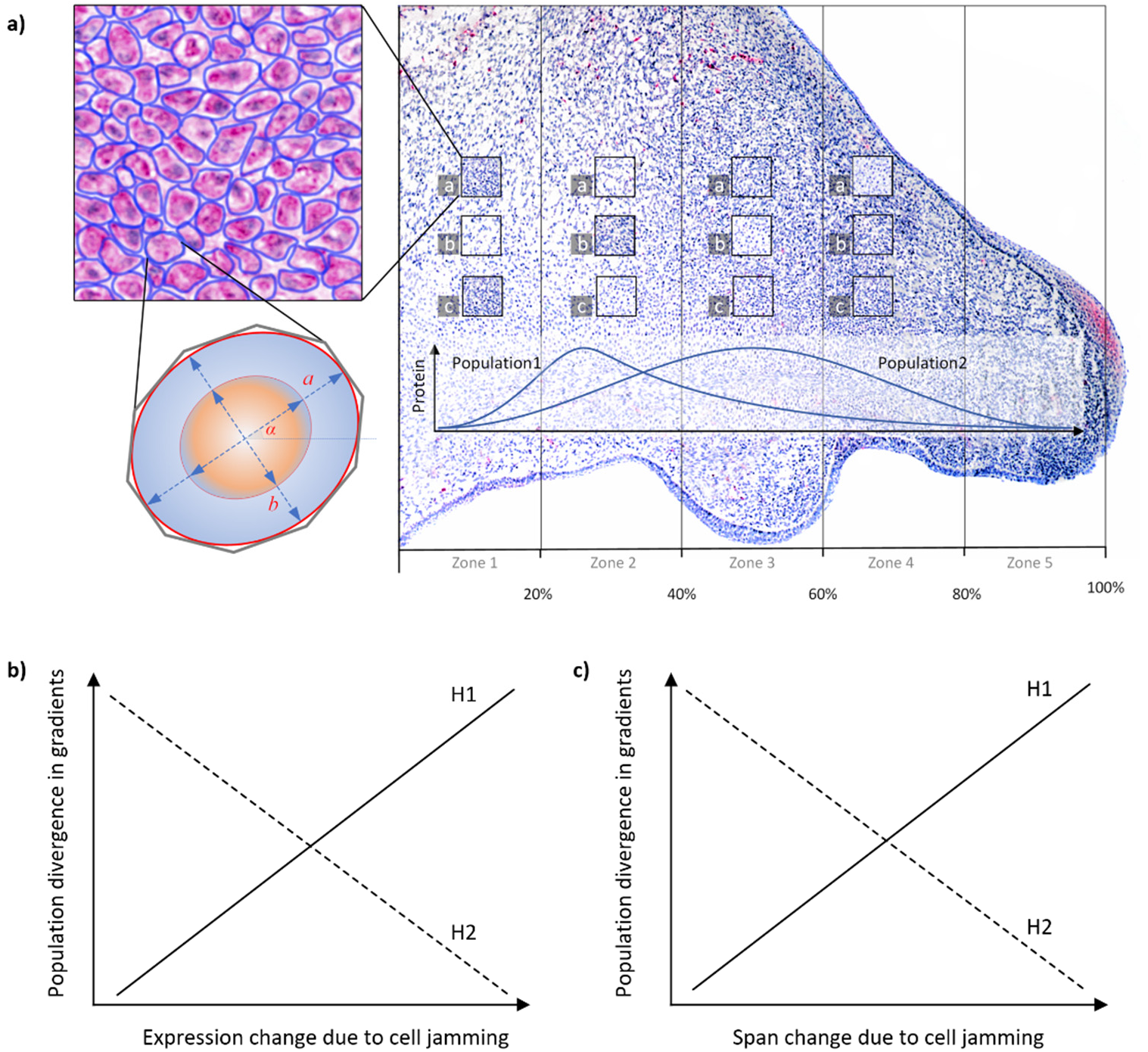
Cell jamming and tissue-level compartmentalization in early ontogeny. **(a)** Schematics of sampling protocol and areas of interest and zones from each beak midline section (see Methods). **(b-c)** We tested whether cell-jamming associated protein expression delineates (H1) or buffers (H2) tissue compartmentalization. (b) Mechanosensing proteins can facilitate tissue compartmentalization by delineating directions of population divergence in protein gradients (measured as Population by Zone interaction, Fig. 3c) through local effects (such as changes in protein connectivity). (c) Cell-jamming induced changes in protein expression can affect anterior-posterior protein gradients priming population divergence in these gradients. Alternatively (H2), the mechanosensing proteins can counteract tissue compartmentalization associated with cell jamming either (b) locally, by accommodating stochastic variation in protein expression caused by jamming or (c) globally by homogenizing spatial reach of protein across tissues.

## 2. Methods

### 2.1 Data collection and sample sizes

Twenty-two study populations of the house finch (*Haemorhous mexicanus*) were grouped into five geographic groups based on their genetic similarity [Supplementary Table S1; 38]. Freshly laid eggs were numbered and either allowed to develop in nests with direct monitoring of incubation onset [39] or placed in portable incubators programmed to house finch incubation parameters [40]. When embryos (*n* = 352) reached the required developmental stage, eggs were opened, and embryos were placed in a petri dish with PBS to be photographed under a Series 80 microscope at 10x magnification. Embryological stage was confirmed [41] and embryos were stored in PAXgene Tissue System (PreAnalytics) that allows simultaneous storage of RNA, DNA and tissues in the field [42–44]. Upon return to the lab, beaks were separated, cryosectioned at 8 µm (within one-cell-thickness, see below) and stored at −80°C. Thirteen sections per individual were obtained at beak midline: one section was stained with Alcian blue hematoxylin and eosin (H&E; U. Rochester MC) to delineate the histological and anatomical area of interest (AOI), twelve were used in immunohistochemical (IHC) analysis of eight proteins described below (Supplementary Figs. S2-S9). Sections were photographed at 600 dpi resolution with a DP74 camera using an Olympus BX43F microscope under x4, x20 and x40 magnifications. To increase statistical power for some tests, twelve developmental stages were combined into three groups: HH25-31, HH32-33, and HH34-36 (Supplementary Table S1), but for each stage only pre-condensation or outside of condensation cells were sampled (see below).

### 2.2 Cell measurements and method validation

We measured shape, density, and alignment of 454,988 mesenchymal cells in 7,408 20x fields of view (60-300 cells in each, hereafter cell groups) distributed across five zones of the upper beak of 352 embryos (Fig. 1a, Supplementary Table S1, Appendix S1). Because our focus was behavior of homogeneous cells, we only sampled cells either prior to formation of condensations, or, at later stages, only cells outside of forming condensations or differentiating tissues. Cell measurements and IHC expression were conducted within an AOI of the upper beak that was delineated based on landmarks homologous across developmental stages and confirmed with H&E histological staining [45]. Briefly, the upper beak AOI was from the point of inflection of the upper beak, to the lower outer tissue of the brain/eye to the inner edge of the mouth, just past the palatine process, but not including the palatine process, to the tip of the beak, along the inner edge of the egg tooth and back to the point of upper beak inflection.

Each AOI was divided into five equal zones (Fig. 1a). Three non-overlapping fields of view of 71μm x 71μm (a, b, c) were randomly selected within each zone under x40 and imaged under 600 dpi. Only mesenchymal cells were sampled, and care was taken to avoid tissue gaps and folds, condensations, epithelial boundaries, and capillaries. The photographs were exported into ImageJ [46], retaining their calibration, and converted to black and white. Custom script thresholds [45] were applied by subtracting the image from its background, enhancing boundaries, and applying an intensity threshold.

We validated our methods in two ways. First, we manually measured cells and nuclei to assess the accuracy of an automated workflow [45] and to verify that nuclei size was a good proxy of cell size. We selected 50 individuals with membranal β-catenin expression (IHC methods, below) across nine developmental stages and imaged the three replica fields of view mentioned above under 40x magnification. Between 10 and 30 cells were sampled from each replica. Photographs were imported into ImageJ and the polygon selection tool was used to manually trace the nucleus and measure area (μm^2^), centroid coordinates, perimeter (μm), major and minor axes of best fitting ellipse, angle of major axis, circularity, and feret length (the longest distance between any two points in the nucleus; Fig. 1a). The same measurements were then repeated for the entire cell. A total of *n* =1,020 cells were measured for this validation. High and repeatable correlations between cell and nucleus area (Supplementary Fig. S11) and shape measures (min feret is shown, Supplementary Fig. S12) confirmed that nucleus size can be used as a proxy measurement of cell size [47–49] and that deformation of mechanosensitive nucleus is proportional and causally linked to the cell jamming transition [50–54]. This enabled us to extend cell morphology measures to IHC assays with only nuclear expression and counterstaining. Second, in the automated workflow [45], three replicas were obtained in each of the five zones as above; the resulting 71μm x 71μm fields of view were photographed under x20 and exported to ImageJ where a custom script recorded nucleus area (μm^2^), centroid coordinates, perimeter (μm), major and minor axis lengths of the best fitting ellipse, aspect ratio, and feret length, angle and coordinates (Fig. 1a). Cells that were overlapping, dividing, or damaged during sectioning were excluded, resulting in complete measurement data for 454,988 cells in 7,408 cell groups (Appendix S1).

### 2.3 Inferring cell jamming state from cell measurements

Variation in cell shape in cell aggregation determines the distribution of contact lengths among neighboring cells and is a sufficient predictor of the jamming transition in groups where all cells experience similar internal and external forces [55], such as in cell monolayers [10, 11]. In most 3D developing tissues that undergo rapid expansions with collective cell movements and rearrangements, such as the mesenchymal cells studied here, neighboring cell groups can differ in fluidity [13, 25], and thus exert and experience variable external forces from other cell groups. In these cases, additional cell metrics, such as cell density and cell alignment [56], might be required to infer onset of jamming.

However even in 3D cell systems, where other cells are treated as a medium (instead of a substrate as in monolayers), the relationship between cell shape and the onset of jamming is fundamentally related to critical values of the length of contacts and contact distribution among deformable cells [57] and corresponding changes in accumulation of compression stress [58]. The main complication for inferring cell mechanical state from geometry alone in such systems comes not from geometry, but from a cell’s ability to actively change their shape, elasticity and other behaviors and thus direct their own jamming and unjamming transitions. Further, even in purely physical systems critical shape values associated with jamming depend not only on dimensionality of the system, but also on context. For example, ellipsoid particles have higher thresholds to the jamming transition than round particles, that jam easier, yet alignment of ellipsoid particles brings them closer to the jamming transition [59]. Broadly however, the same principles apply as indicated by 1) universality of the relationship between shape and its variability in jamming transitions and 2) broad similarity in properties of jamming transitions in physical and biological systems and in monolayers and 3D tissues [57]. Furthermore, metrics utilized by the studies of jamming in 3D tissues, such as critical values of the index of asphericity (p^2^/4πa) [57] are geometrically related to the critical values of the cell shape index (*p_o_*) that we used here in our analysis of cells on 2D IHC slides. Thus, our combination of measures can be used to infer jammed and unjammed states of cell groups during development.

### 2.4 Classification of cell jamming/unjamming

We classified cell jamming for each cell group (in a, b or c ; Fig. 1a) on the one-cell thick section by applying five metrics indicative of jamming transitions and calculating agreement among these metrics. First, we calculated the cell shape index 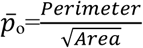 and used *p*\*=3.81 as a critical value below which cells are jammed, such that any cell rearrangements require cell shape changes [10] (minimum value of *p* is 3.54 at which a cell is a perfect circle). Second, to specifically examine the effect of shape elongation, we calculated cell aspect ratio 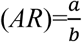 and its variability (standard derivation) and used s.d. (*AR*) =0.5 (corresponding to *AR*≈1.6 in our sample) as a critical value below which cells are increasingly uniform and round and are jammed. Third, we calculated the standard deviation of cell polarization in cell groups (angle *α* in Fig. 1a) and used s.d. (*α*)=0.5 as a critical value below which cells within a group are aligned. Fourth, we calculated total area occupied by cells within the 71x 71μm area and used the critical value of 95% as a cutoff below above which cells are jammed. Fifth, we calculated *AR* for all cells within a group and calculated ratio of cells with *AR*<1.6 to the number of all cells within a group and used critical value of 70% above which cell group was considered “jammed”. We then compared agreement scores (Cohen’s Kappa) across all trios, quartets, and quintets of these metrics. We found that the combination of cell shape index (*p*_o_), variation in cell alignment (s.d. (*α*)), and variation in aspect ratio (s.d. (*AR*)) showed the highest agreement among all the metric combinations (Cohen’s K (overall) = 0.46 (95% CLI: 0.39-0.53), McNemar’s S = 123.05, *P* < 0.001). Agreement on “jammed” classification among these three indices was higher than the agreement on “unjammed” classification (Cohen’s K = 0.53 (0.48-0.58) vs Cohen’s K = 0.39 (0.34-0.44)).

### 2.5 Immunohistochemistry

Sections were blocked with an endogenous peroxidase (2% H_2_O_2_) for 20 minutes, washed in tris-buffered saline (TBS), blocked with 2% bovine serum albumin (BSA) plus 5% normal goat serum (NGS) in TBS for 1 hour. Avidin-Biotin blocking kit (Abcam) was used per manufacturer’s instructions before applying antibodies. Samples were incubated for 18 hrs with a primary antibody at 4°C, washed in TBS, incubated at room temperature for 1 hour with a secondary antibody, and washed again with TBS. Appropriate dilution of the primary antibody was determined by running preliminary trials with serial dilutions. Primary antibodies included anti-β-catenin (610153, 1:16,000, BD Transduction Laboratories), anti-CaM (sc-137079, 1:15, Santa Cruz Biotechnology), anti-Wnt4 (ab91226, 1:800; Abcam), anti-TGFβ2 (ab36495, 1:800, Abcam), anti-Bmp4 (ab118867, 1:100, Abcam), anti-Ihh (ab184624, 1:100, Abcam), anti-Dkk3 (ab214360, 1:100, Abcam), and anti-FGF8 (89550, 1:50, Abcam). Three different secondary antibodies (Biotinylated Goat Anti-Mouse IgG, BP-9200; Biotinylated Goat Anti-Rabbit IgG, BP-9100; Biotinylated Goat Anti-Rat IgG, all 1:200, Vector Labs) were used depending on the host primary antibodies. Negative controls were incubated with TBS instead of primary antibodies. During optimization of assays, controls were also run with just the secondary antibody to verify there was no non-specific binding. Validation assays, including isotype controls, for specificity of stains is shown in Supplementary Figures S3-S10. Reactions were visualized with either diaminobenzidine (DAB, Elite ABC HRP Kit, PK-6100, Vector Labs) or Vector Red Alkaline Phosphatase substrate and Vectastain ABC-AP Kit (AK-5000, Vector Labs). Hematoxylin was used to counterstain nuclei. Each slide contained four sections, and the three slides (12 sections per embryo) were run with the following grouped antibodies: i) β-catenin, Fgf8, Tgfβ2 and no primary control, ii) Bmp4, Wnt4, Ihh and no primary control, and iii) Dkk3 and no primary control, and CaM and no primary control (Appendix S2). Suitable upper beak midline sections with expressed proteins (*n* = 5,044) were imaged and named according to embryo ID, protein, developmental stage, and IHC run to enable automated processing as described in Lee et al. [45]. Our processing protocol randomized assignment of sections from different populations and stages to IHC runs.

### 2.6 Quantification of expression and spatial reach of proteins

We used a custom script to measure protein expression across AOIs [45]. Briefly, the script partitioned AOIs into five equal zones (20% of the anterior-posterior span) without image rotation to prevent data distortion, converted the image to black and white, applied a specific threshold for each protein expression and outputted the area of expressed, unexpressed and total tissues for each zone and for the entire AOI (details in [45]). Spatial reach (also known as “correlational span” in literature, see below) was a percentage of anterior-posterior span of five zones of upper beak (in 20% increments) over which the expression of protein was significantly correlated [after 22]. Only adjacent zones were included in calculations. Zones were considered “unjammed” if at least two of the three cell groups (a, b, c) were in the unjammed state, and “jammed” if at least two cell groups were in the jammed state. All zones across spatial reach had to be in the same state to be included in calculations. Supplementary Fig. 10 shows details and examples of calculations.

### 2.7 Statistical analyses

To achieve normal distribution, reduce skewness and stabilize variance, we used the Box-Cox transformation with λ=0.5 for raw data on protein expression, log10 transformation for cell morphology measures and arcsine transformation for cell density proportional measures. We used a mixed-effects model with Restricted Maximum Likelihood to examine the effects of cell jamming state on protein expression. Population and embryo ID were treated as random effects. Significance of Zone was examined as an interaction between Zone and embryo ID. Population divergence in anterior-posterior protein gradients were modeled as Population by Zone interaction. We computed least squared means of protein expression for each cell state (jammed and unjammed) and assessed significance of these means and difference between them with a Sidak adjustment for multiple comparisons (Supplementary Table S4). To directly compare the contribution of both random and fixed effects to variance in protein expression explained by mixed effects models, we compared their contribution to Adjusted R². We first constricted a full model incorporating all fixed effects and their interactions. We then generated a set of reduced models, starting with the intercept-only model to assess baseline variance and sequentially omitting each predictor in other models. For each model, the residual variance estimates were extracted, and Adjusted R² was computed as: Adj R² = 1 - (Residual Variance (Reduced Model) / Residual Variance (Null Model)). The contribution of each predictor was then calculated as the difference in Adjusted R² between the full model and each reduced model: Percent Contribution = ((Adj R²_Full - Adj R²_Reduced) / max(Adj R²_Full, 0.0001)) × 100. This approach allowed us to quantify the independent explanatory power of each predictor while accounting for shared variance in the model. We then ranked these contributions to Adj R² for each predictor for Fig. 2 (Supplementary Table S4).

**Fig. 2.**
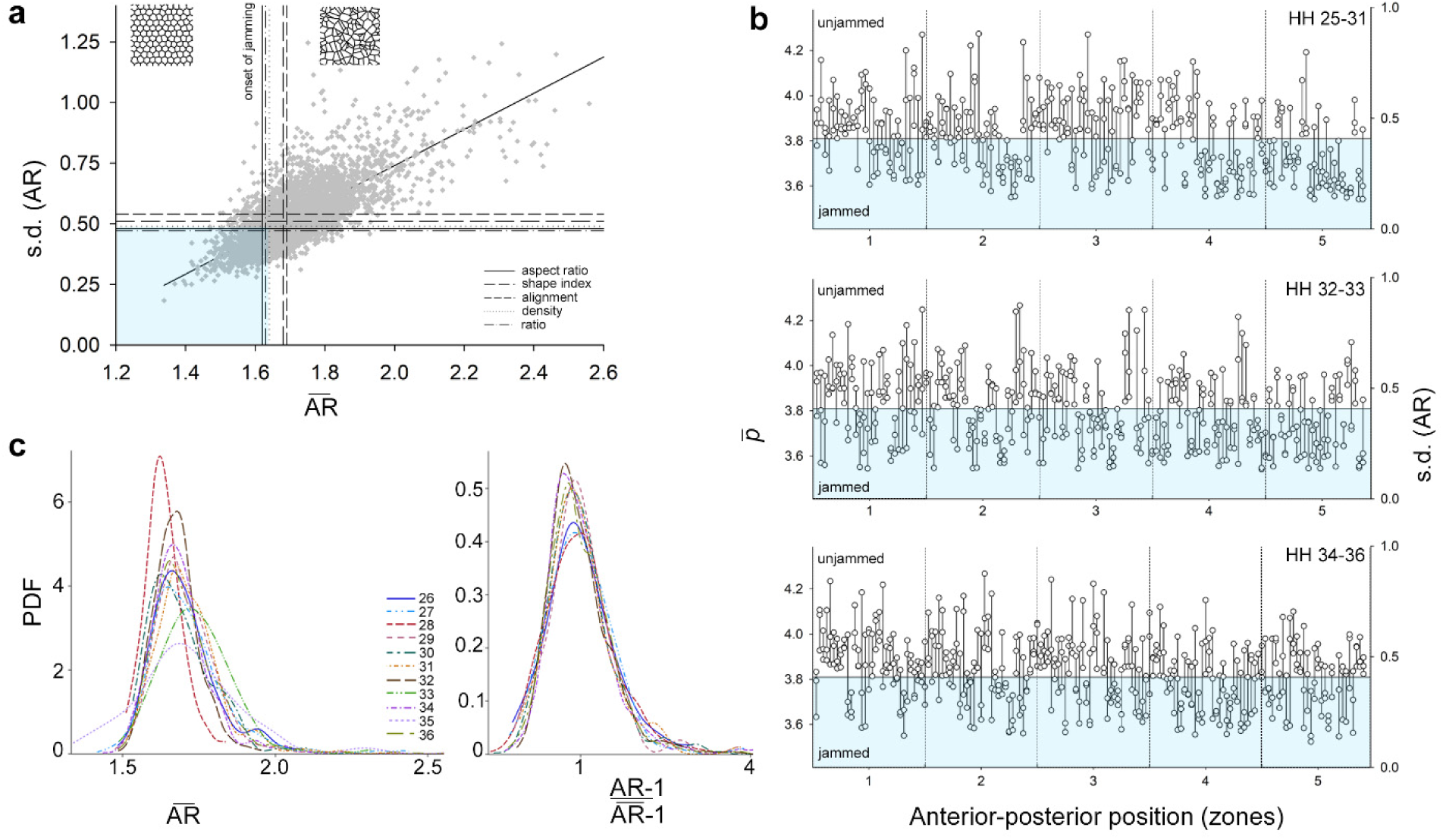
Cell jamming transitions across biological contexts. **(a)** Relationship between the average cell aspect ratio (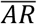) and its variability (standard deviation of *AR*) in each cell group, and comparison of five metrics predicting onset of cell jamming (see text). Shaded rectangle outlines values of (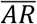) and s.d. (*AR*) that correspond to the agreement between all five metrics in classification of jammed state. **(b)** Distribution of jammed and unjammed groups of cells at the same anterior-posterior position and across developmental stages (HH25-31 upper, HH 32-33 middle, HH 34-36 lower; Supplementary Table S3). Lines connect groups of cells (dots, 60-300 cells in each; a, b, c viewframes in Fig. 1a) of the same sample. Shaded area corresponds to the onset of jamming. Left axis shows critical values of average cell shape index (*p̅*) and right axis – s.d. (*AR*) for reference. **(c)** Probability density functions (PDF) for cell 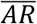 across developmental stages (left graph, populations combined, Supplemental Fig. S1b shows each population) collapse into a common distribution upon AR rescaling (right graph).

## 3. Results

### 3.1 Cell jamming is linked to the scaling of cell shape

We find that cell shape (measured as aspect ratio, *AR*) and its variability (measured as s.d.) are linked: as cells in a group become more isotropic (due to more similar and equally distributed contacts with neighboring cells on all sides), they become less variable (*b*_ST_ = 0.76, *t* = 39.66, *P* < 0.0001; Fig. 2a). The slope of the relationship between 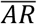 and s.d. (*AR*) did not differ across developmental stages, populations, and proliferation zones (Supplementary Table S2) and likely represents a geometric constraint linking cell shape variability and tissue rigidity, as often observed in both cell monolayers and 3D cell aggregations [11, 54, 60–62]. To examine whether packed groups of increasingly uniform and rounder cells represent a jammed state, we calculated the agreement between five metrics commonly deployed to predict the onset of cell jamming (see Methods). To visualize the agreement between all five metrics, we show critical values associated with the onset of jamming on 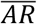 vs s.d. (*AR*) plot (Fig. 2a, *p̅\** = 3.81, s.d. (*α\**) = 0.5, s.d. (*AR\**) = 0.5, density = 95%, ratio = 70%). We find that throughout early development, groups of cells transit between jammed and unjammed states (Fig. 2b). In all populations and developmental stages, groups of jammed cells are local (having more than two jammed cell groups within a grid zone was uncommon) and randomly distributed across the grid (Fig. 2b, Supplementary Table S3). Probability density functions (PDF) of cell 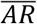 derived for developmental stages, populations, and zones are all unimodal, but differ slightly in distribution (Fig. 2c, Supplementary Fig. S1a). We thus tested the commonality of underlying functions across these biological contexts, by rescaling *AR* for each PDF to mean=1 and min=0. We find that all PDFs of rescaled 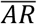 collapsed into a shared density function (Fig. 2c) with minor differences across otherwise widely distinct biological contexts (e.g., all developmental stages in Fig. 2c and all populations and zones in Supplementary Fig. S1b).

Having established that interdependence of cell shape and its variability (Fig. 2ac) underlies jamming transitions across tissue (Fig. 2b), we then asked whether the network of regulatory proteins that are central for beak tissue differentiation [Supplementary Figs. S2-S9, 35, 36] take advantage of this ubiquitous physical process.

### 3.2 Spatial reach of key proteins is associated with cell jamming

We find that within-zone expression of Dkk3, Bmp4, Ihh and Tgfβ2 decreases when tissues are in the jammed state, whereas the expression of Fgf8, CaM, Wnt4 and β-catenin increases (Fig. 3a). We then examine changes in the spatial correlation of protein expression along the anterior-posterior axis of the upper beak in relation to jamming states of zones (Fig. 3b, Supplementary Fig. S10). Following previous studies, we call this metric *spatial reach*, it is also called decay length, correlational span, or correlational length in the literature [e.g., 22, 31]. We find that under the jammed state, the expression of β-catenin, Dkk3, Tgfβ2 become more spatially restricted, whereas the expression of CaM, Fgf8 and Wnt4 became more spread-out and uniform (Fig. 3b). With the exception of β-catenin where a local increase in expression is associated with more fragmented global distribution along the jaw (Fig. 3a,b), all proteins show concordant changes in the direction and magnitude of cell-jamming associated expression and its spatial reach across jaw (Fig. 3e). We next examine divergence in protein gradients in early beak primordia across recently established populations and contribution of protein sensitivity to local cell jamming transitions to this divergence.

**Fig. 3.**
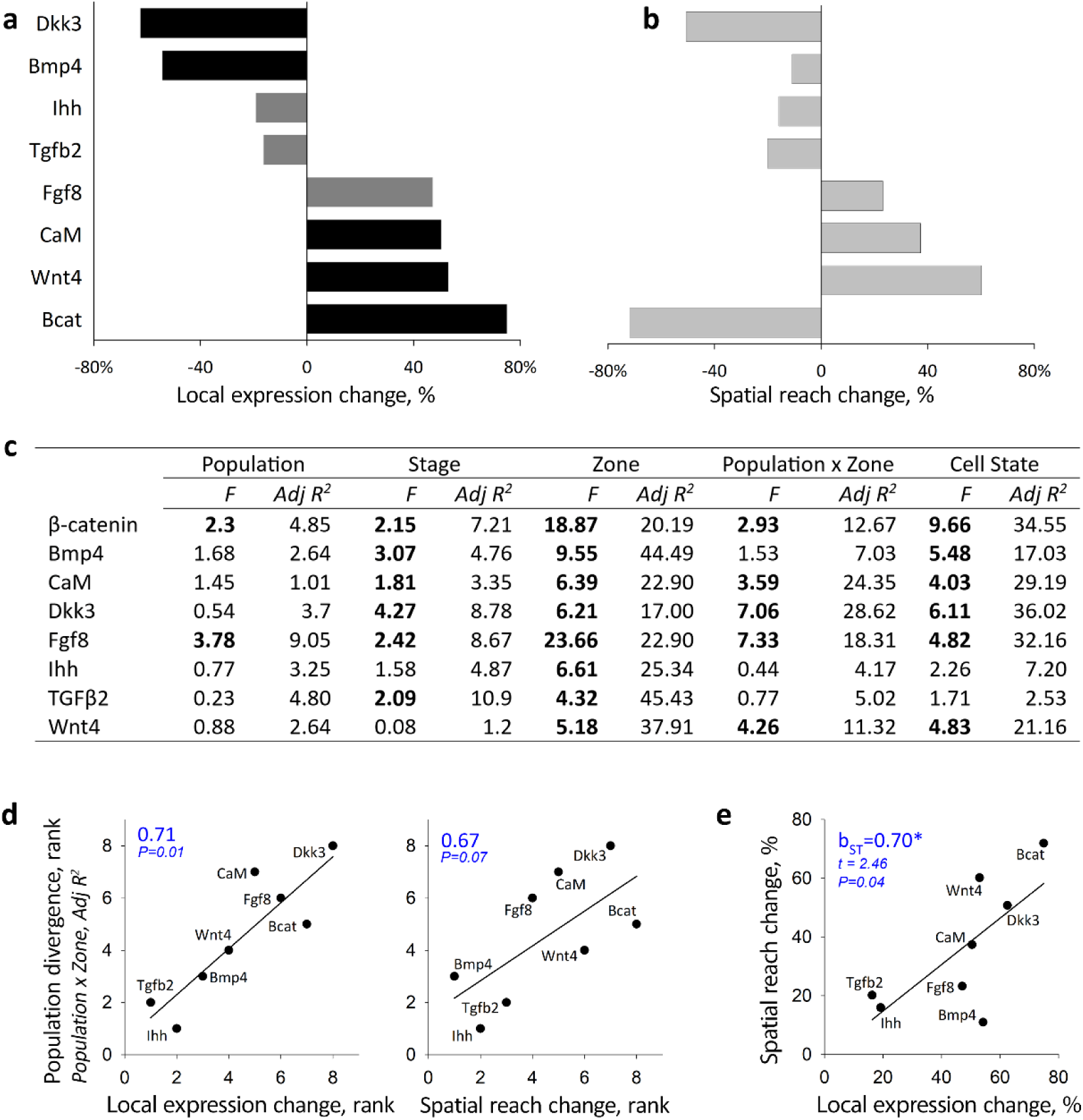
Protein expression and spatial reach changes induced by cell-jamming prime regulatory population divergence. **(a)** Local (within zone) changes in protein expression between jammed and unjammed groups of cells. Bars show relative change (%) between the least squared means (mixed effects model in *c*) for jammed and unjammed cell states. Black bars: *P* < 0.01, dark gray bars: *P* <0.05 (data in Supplementary Table S4). **(b)** Change in spatial reach of protein (a percentage of total continuous anterior-posterior length over which protein expression is correlated; data from Supplementary Fig. S11 that shows details of calculations and Supplementary Table S4) when spanning jammed vs unjammed groups of cells. **(c)** Mixed effect model estimating the effects of population, developmental stage, zone, population divergence in protein gradient (Population x Zone) and cell state (jammed vs unjammed) on protein expression. Shown are *F*-values (bold shows *P* < 0.05 after within-model Sidak adjustment) and variance contribution (in %) to Adjusted R^2^. **(d)** Cell-jamming induced change in within-zone protein expression (left) and its effect on spatial reach of proteins (right) predict population divergence in protein gradients. Ranks are Adj R2 variance contributions from (c), Spearman correlation coefficients are shown. **(e)** Cell-jamming induced changes in within-zone protein expression predicts changes in spatial reach of the protein (absolute values, *b*_ST_ is standardized regression coefficient, data from Supplementary Table S4).

### 3.3 Cell jamming primes divergence in tissue compartmentalization

Anterior-posterior gradients of protein expression differ among recently diverged populations (Figs. 3c, 4). For example, Dkk3 expression at HH25-31 is the highest in posterior proliferation zones in eastern and northwestern Montana populations, but in the middle zones in central and northern Montana. Similarly, the anterior-posterior gradient of Tgfβ2 and CaM during HH25-31 diverges among populations (Fig. 4). Divergence in anterior-posterior gradients (modeled as Population by Zone interaction in the mixed-effects model) is the strongest in CaM, Dkk3, and Fgf8, whereas Bmp4, Ihh and Tgfβ2 gradients are similar across populations (Fig. 3c). We then examine the association between protein expression and spatial spread induced by cell jamming and population divergence in protein gradients (Fig. 1b,c). We find that protein responsiveness to cell jamming – both in local expression and in spatial reach – predicts population divergence in protein gradients (Fig. 3d), supporting the hypotheses that cell jamming facilitates regulatory compartmentalization and divergence at the level of tissues (Fig. 1b,c).

**Fig. 4.**
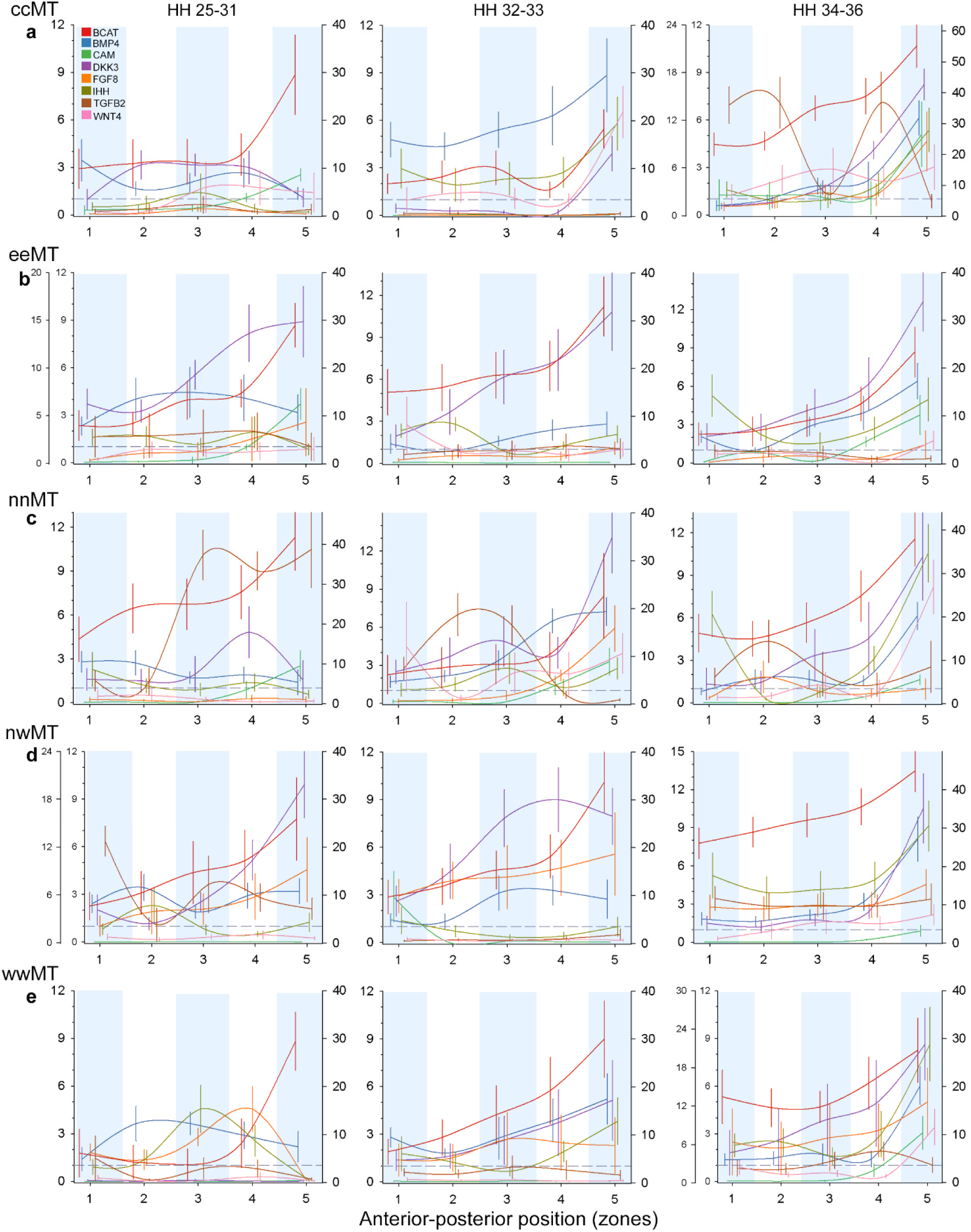
Population-specific gradients of protein expression across anterior-posterior position and developmental stages. Shown are means for each zone ± SE. Developmental stages arranged in columns. The left-most ordinate axis (on graphs with two left axes) shows the expression of Dkk3, the right ordinate axis shows the expression of β-catenin. Expression of all other proteins is shown by the left ordinate axis. Gray dashed line shows the threshold of IHC detectability. Populations (Supplementary Table S1): **(a)** ccMT – central Montana, **(b)** eeMT – eastern Montana, **(c)** nnMT – northern Montana, **(d)** nwMT – northwestern Montana, and **(e)** wwMT – western Montana.

## 4. Discussion

We found that transient and local jamming transitions in pre-condensation mesenchymal cells, linked to the scaling of cell shape, were associated with the spatial reach of protein expression across the mesenchyme of the upper beak in a pattern that is mirrored in the regulatory divergence among populations. Close association between the activity of morphogens, transcription factors and other regulatory proteins and aspects of the cell jamming transition [13, 25–27, 35, 63] have three main implications. First, these findings show how local transitions in fields of homogeneous cells can accomplish coordinated and region-specific cell behaviors over longer spatial scales, leading to progressive compartmentalization of tissues – a prerequisite for developmental specification and complexity [e.g., 64, 65]. Second, the linkage between regulatory proteins’ expression and material properties of cell groups is well placed for evolutionary modifications of morphogenesis. This is because the entanglement of path-independent (i.e., ahistoric) physical processes and highly contingent (i.e., path-dependent) genetic and biochemical processes imposes history-dependence onto otherwise simple-path processes or resets highly continuous biological processes. In development, such integration allows for cycles of modification and stabilization that are necessary for differentiation and accommodation of stochastic noise [11, 13, 17, 66]. In evolution, dynamic integration of processes with distinct history-dependence allows developing systems to adjust and evolve without losing robustness [67, 68] and thus transit between adaptations. Third, the minimal rules that govern individual cells modify the material state of tissues and are ultimately detectable in microevolutionary population divergence. What are the mechanisms behind these scale-transcending effects?

If the transient groups of jammed mesenchyme cells studied here are developmental precursors of “cell condensations”– fundamental units of cellular differentiation and morphological diversification in avian beaks [69–73] – then this might explain the mesoscale effects of cell jamming transitions and their imprint on the patterns of population divergence (Fig. 3). Beak condensations arise when the neural crest cells, induced from the developing neural tube, migrate in distinct streams (depending on the site and time of their induction) to different locations in the upper beak where, upon interaction with the beak epithelium, they form condensations [74–77]. We envision three general scenarios by which transient cycles of jamming in mesenchymal cells can be relevant to the origin of condensations (Fig. 5). First, jamming can synchronize cell cycles in groups of mesenchymal cells (Fig. 5a), either by erasing differences accumulated during migration and thus priming these cells for a transition to a condensation (scenario *i* in Fig. 5a) or by synchronizing and amplifying signaling arising from cell-cell interactions within jammed cell groups [Fig. 5a, ii;78, 79]. Second, uniformly distributed small islands of jammed cells (Fig. 1b, Supplementary Table S3) can create larger scale standing waves of morphogen signaling that prepattern the sites of future condensations by priming cells within these areas and affecting cell migration [Fig 5b; 80]. Cell jamming transitions and associated cell-scale stresses can also delineate the boundaries of future condensations by “melting” jammed tissues and channeling cell migration and accumulation or by sorting cells according to their jammed state [63, 81–85]. Finally, condensations can originate when jammed cell groups exceed a critical size that prevents their dissociation through either mechanical or signaling mechanisms [Fig. 5c; 63].

**Fig. 5.**
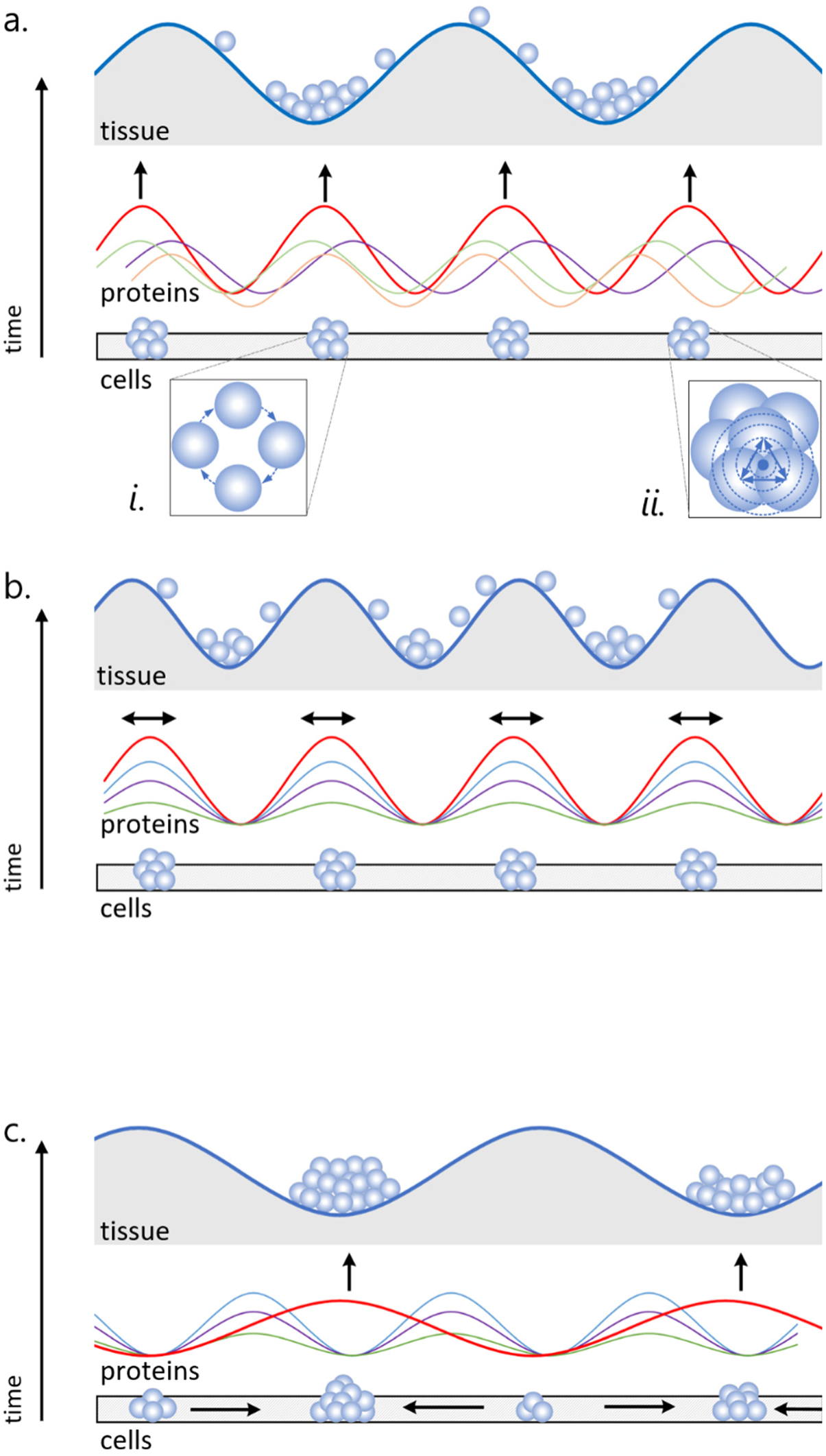
Linking cell jamming transitions and the origin of cell condensations. Proposed mechanisms by which transient groups of jammed cells (low: “cell level”) can be developmental precursors of persistent cell condensations (upper: “tissue level”) that arise at subsequent developmental stages at the interface of epithelium and mesenchyme. (**a)** Signaling (“proteins”) associated with dynamic cell-cell interactions during jamming transitions prepattern future condensations. *i*) jamming resets and synchronizes cell cycle variability among mesenchymal cells arising from distinct origins or arriving by different migration routes, priming them for condensation transition, *ii)* jamming amplifies signaling in the jammed cell group. (**b)** Groups of jammed cells form boundaries for future condensations either by juxtaposition of protein signaling gradients or through cell adhesion and sorting. **(c)** Condensations originate from jammed cell groups above critical size or duration of existence either through amplified signaling or through local cell adhesion.

More broadly, developing systems reconcile two seemingly opposite properties – they are robust to perturbations and adaptively flexible at the same time [20, 68, 86]. In a systems biology view, these two requirements correspond to the properties of, respectively, an ordered phase (that enables stability and small step incremental improvements) and a disordered phase (that enables large steps in the search for novel solutions). This has led to the postulate that evolution keeps biological systems near a critical transition – just inside the ordered phase, but able to rapidly transit to a disordered phase [24, 87–89]. Our finding of notable non-linearity of the effects of cell mechanical state on the spatial reach of regulatory proteins, where small changes in the size and shape of individual cells are linked with long-range correlations of protein expression, and thus large expansions of developmental scale (Fig. 3), is characteristic of such a critical transition.

Specifically, our results corroborate the idea that components of development differ in their distance to critical transitions and that these dynamic changes in criticality can reconcile robustness and evolvability of development [8, 23, 90]. First, whereas the transition between levels of organization (e.g., cells and tissues) might be close to criticality, stability is favored within each level as is evident in the remarkably homogeneous distribution of cell shapes and jamming states (Fig. 2, Supplementary Fig. S1, Supplementary Table S3). Second, the interchangeability of factors that lead to a jamming transition virtually guarantees that adjacent areas will be asynchronous in their jamming cycles, and thus, their distance to critical transitions. This will lead to their compartmentalization and differentiation despite uniformity of their elements. Third, some aspects of the regulatory protein network that are closely integrated with material properties of tissues (called dynamic modules in other studies) might be close to critical transitions while others remain stable [23, 67, 90], thereby channeling developmental variation and evolutionary divergence. Our finding of the variable association of protein expression and spatial reach with cell jamming is consistent with this idea (Fig. 3, Supplementary Fig. S10).

The latter point raises the question of the possible mechanisms behind variable mechanosensitivity of protein expression in this system. In a companion study, we showed that mechanosensitivity of transcriptional regulators was determined by their intrinsic disorder – disordered proteins mediated jamming transitions through their dosage-dependent binding plasticity [91]. Dosage-dependent binding plasticity of intrinsically disordered proteins can act as a developmental reset mechanism – forming the dynamic modules described above and channeling developmental and genetic variation in development. The mesoscale effect of local jamming transitions on proteins’ spatial reach (Fig. 3b) also raises the possibility of transcellular protein trafficking or the effect of tissue rigidity on morphogen spread as potential mechanisms [92].

Finally, a defining feature of criticality is expanding the spatial reach of effects at each transition, which is also a defining feature of scaling transformations. In this respect, it is notable that the remarkable diversity of avian beak shapes can be explained by simple rescaling along a few developmental axes [33, 93, 94]. Developmentally, scaling transformations are often a product of modification of gradients of diffusible secreted morphogens or growth factors, such as Bmp4 or Tgfβ2 studied here [92]. Integration of material properties of tissues with the propagation of morphogens (Fig. 3), enables precise control of morphogen behavior in relation to tissue size [31, 92]. Thus, local changes in cell growth parameters together with the scale-expansion of these effects (Fig. 2) could well be the ontogenetic origin of scaling transformations, such as those seen in avian beaks [37, 95].

In sum, we find that a combination of cell jamming with mechanosensitivity of key proteins transiently and yet, predictably, resets developmental scale, priming regulatory tissue compartmentalization and uncovering rich opportunities for evolutionary diversifications while preserving developmental integration and robustness.

## Data availability

All data are available in the manuscript and the supplementary materials.

## Acknowledgments

We thank Fatima Bravo, Kaitlyn Gahl, Kathryn Chenard, Chris Seliga, Xander Posner, Robert Hollingsworth, Jakob Abtahi, Omar Puebla, and Ali Shaikh for help with collection and preparation of samples, cryosectioning, histological and molecular assays and computational work, and Neha Varshney and Arhat Abzhanov for generously sharing their expertise in histology and immunohistochemistry.

## Author contributions

AVB designed the project and obtained funding; AVB, CAL, MJG, GAS, CSM, SEB, RAD developed methods, collected data, and analyzed data. AVB wrote original draft, AVB, CAL, MJG, CSM, SEB, RAD reviewed and edited final version.

## Supplementary Materials for

**Fig. S1.**
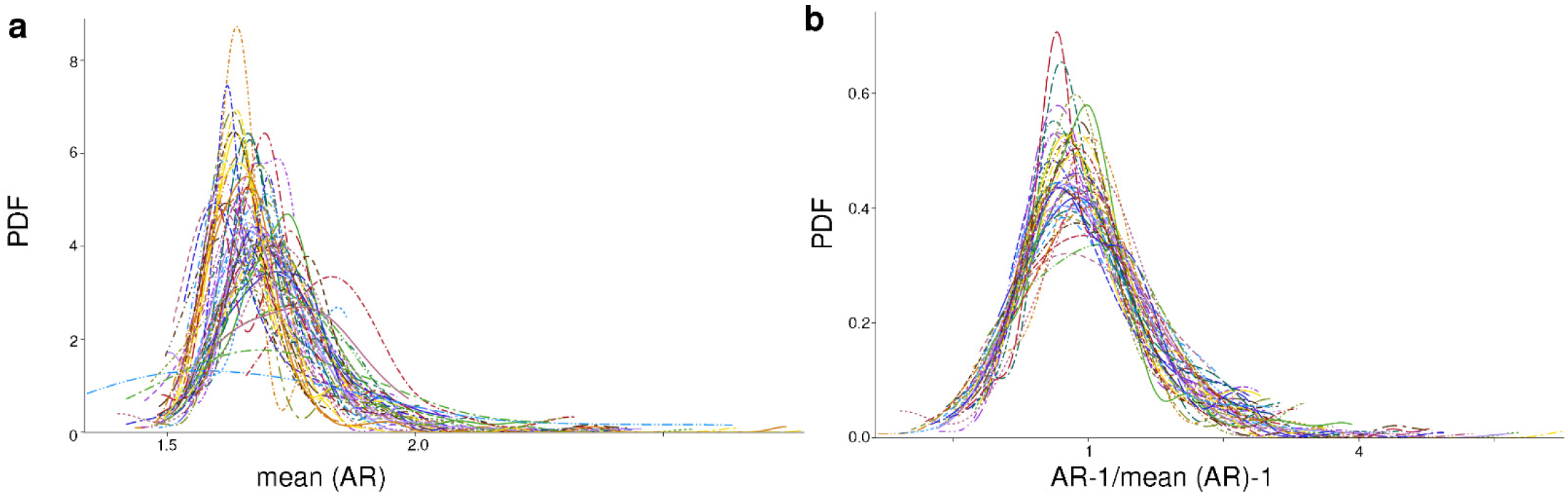
**(a)** Probability density functions (PDF) of mean (AR) of cells (*n* = 454,988) for all zones, populations, and developmental stages – representing the entire variability of cell shape across early ontogeny across all stages and contexts in the dataset (Appendix S1) – collapse into a near-common distribution **(b)** upon rescaling.

**Fig. S2.**
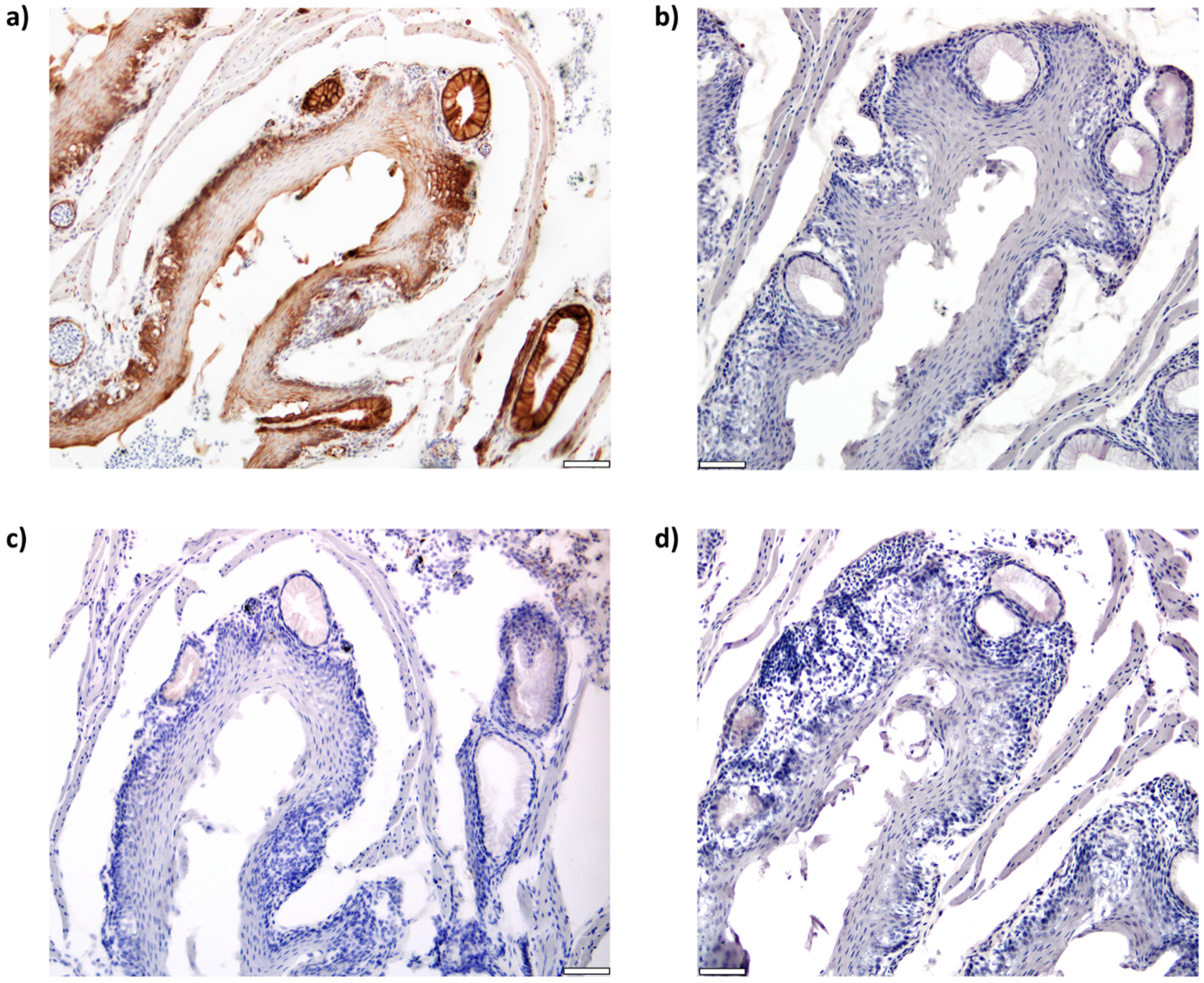
β-catenin expression in zebra finch esophagus with positive and negative controls. **a)** Specific expression observed in epithelial layer and mucous glands and no expression in blood cells. No expression is observed in the **b)** isotype, **c)** no primary, and **d)** no secondary controls, validating specificity of staining. Scale bar is 50 μm.

**Fig. S3.**
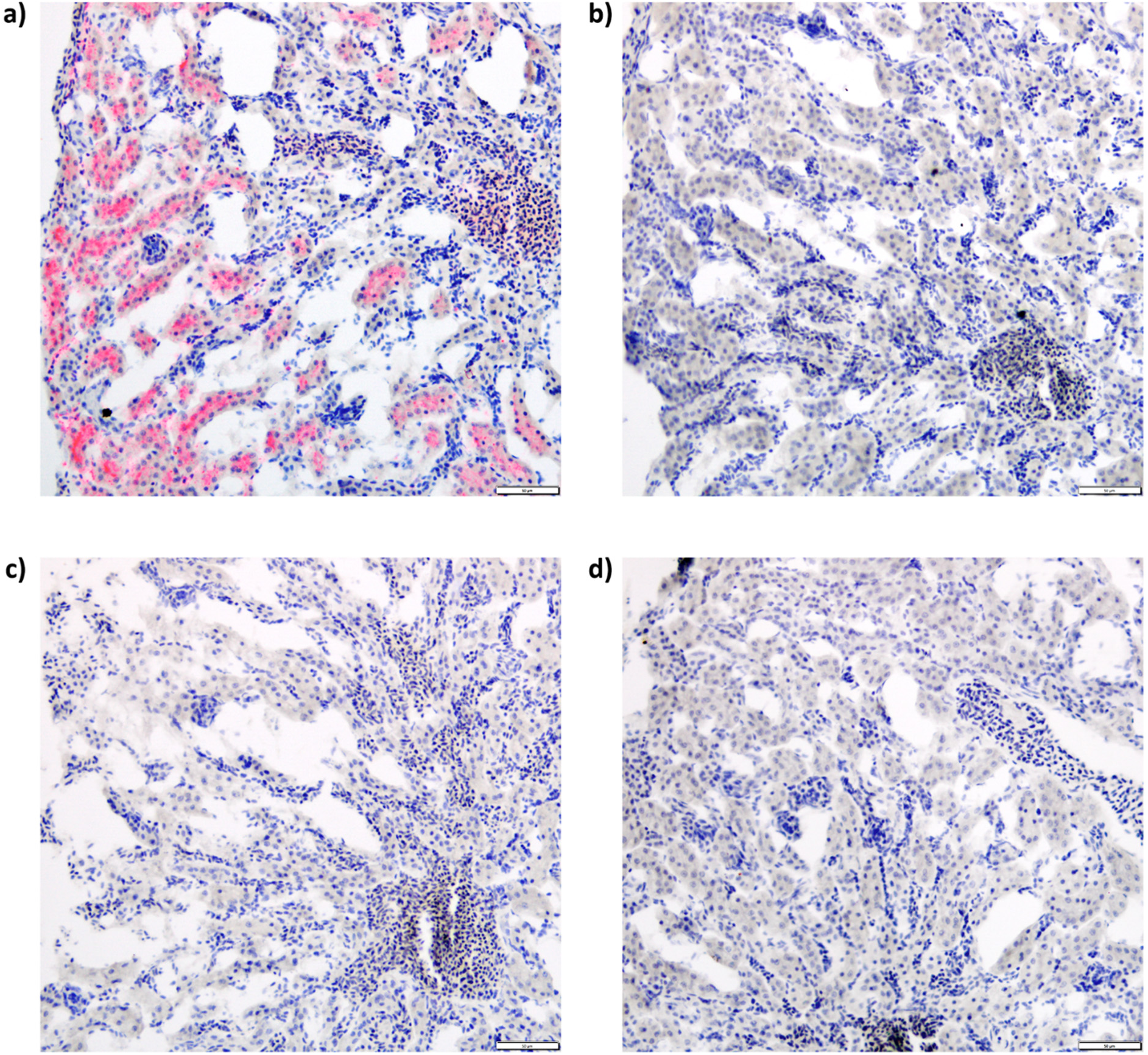
Bmp4 expression in house finch kidney with positive and negative controls. **a)** Specific expression observed in extracellular matrix with negative expression in glomeruli. No expression is observed in the **b)** isotype, **c)** no primary, and **d)** no secondary controls, validating specificity of staining. Scale bar is 50 μm.

**Fig. S4.**
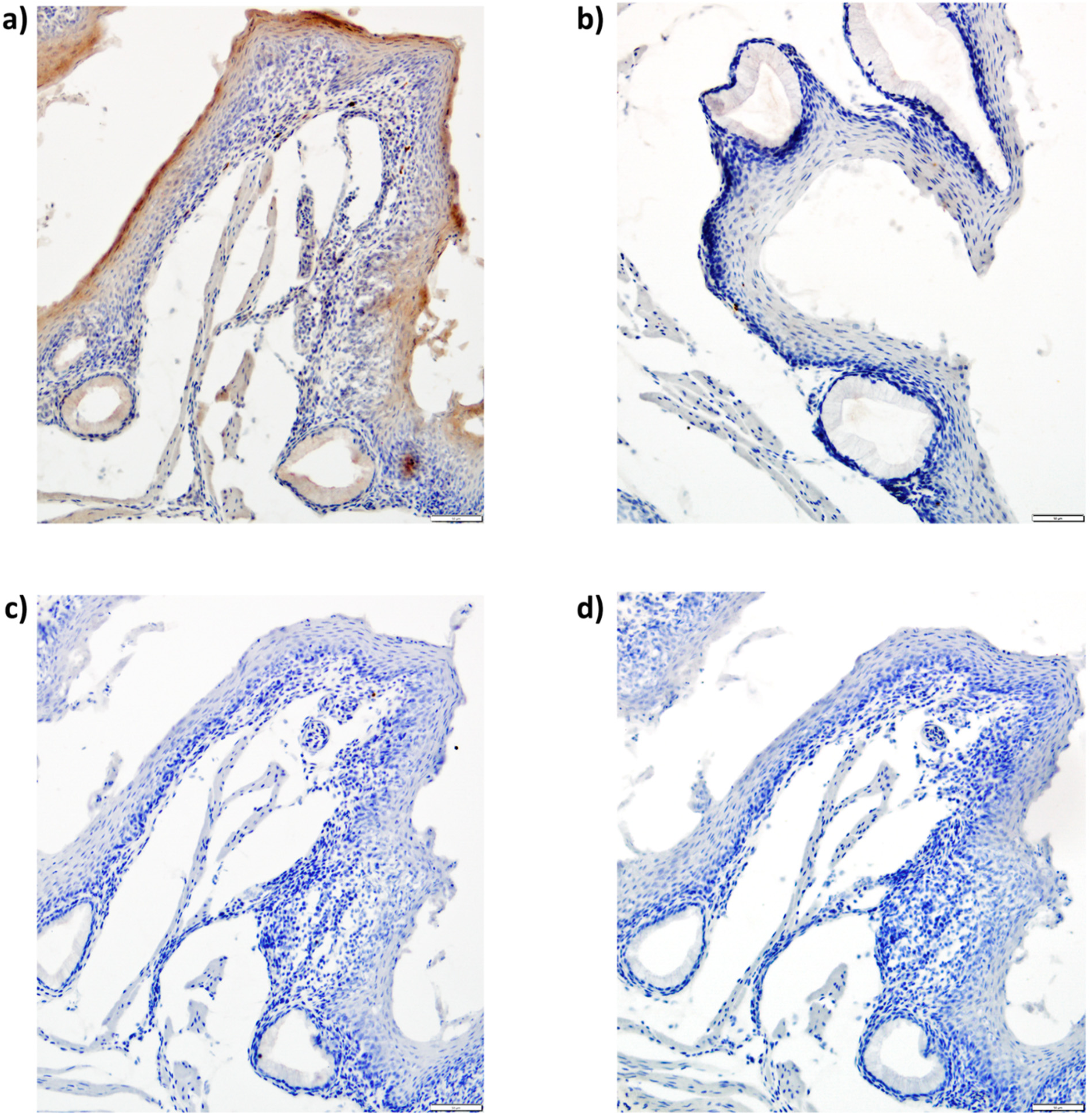
Calmodulin 1 expression in zebra finch esophagus with positive and negative controls. **a)** Specific expression observed in epithelial layer, some light expression in glands, and no expression in submucosa and muscle. No expression is observed in the **b)** isotype, **c)** no primary, and **d)** no secondary controls, validating specificity of staining. Scale bar is 50 μm.

**Fig. S5.**
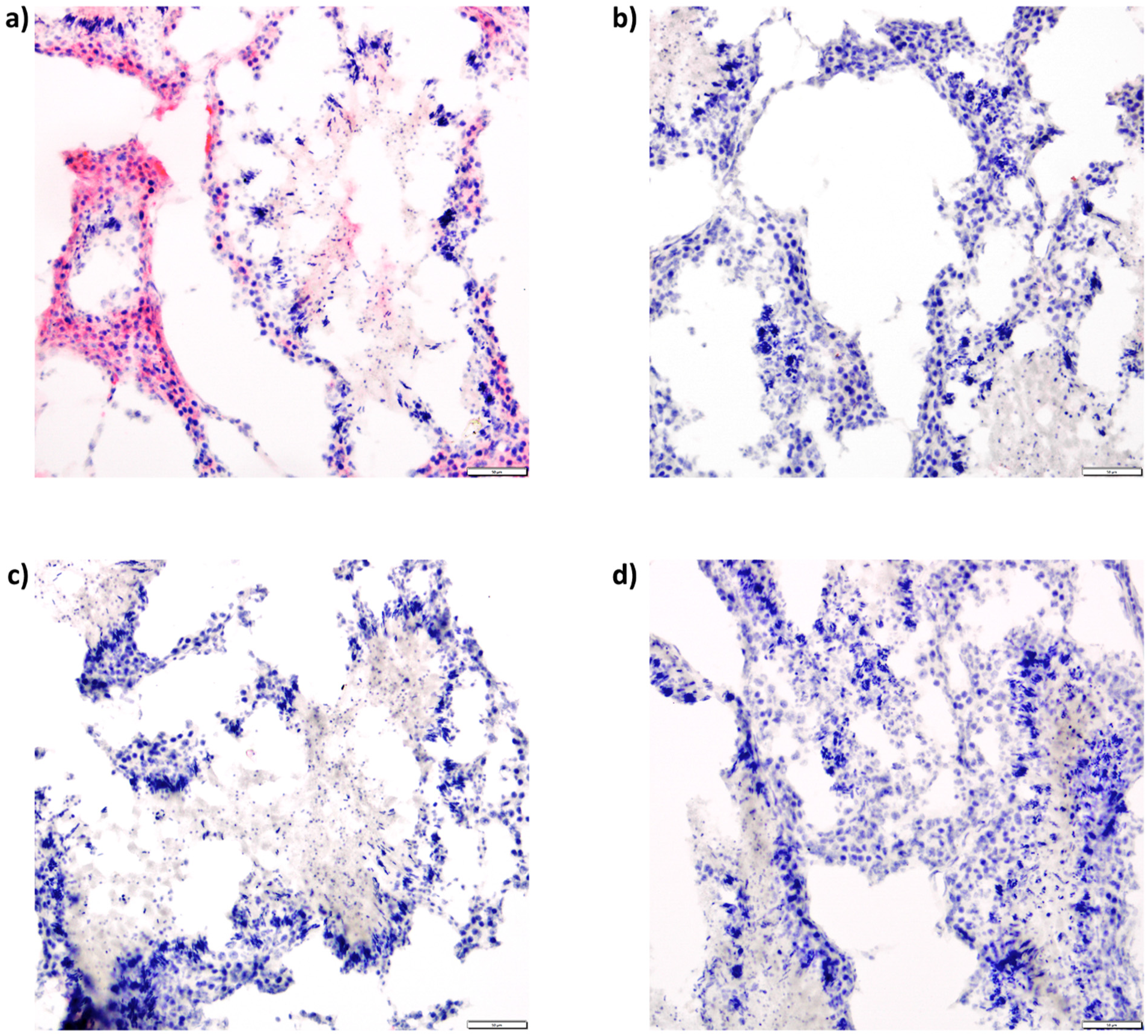
Dkk3 expression in zebra finch testes with positive and negative controls. **a)** Specific expression observed in the interstitial cells and spermatogonia and no expression in spermatocytes. No expression was observed in the **b)** isotype, **c)** no primary, and **d)** no secondary controls, validating that staining is specific. Scale bar is 50 μm.

**Fig. S6.**
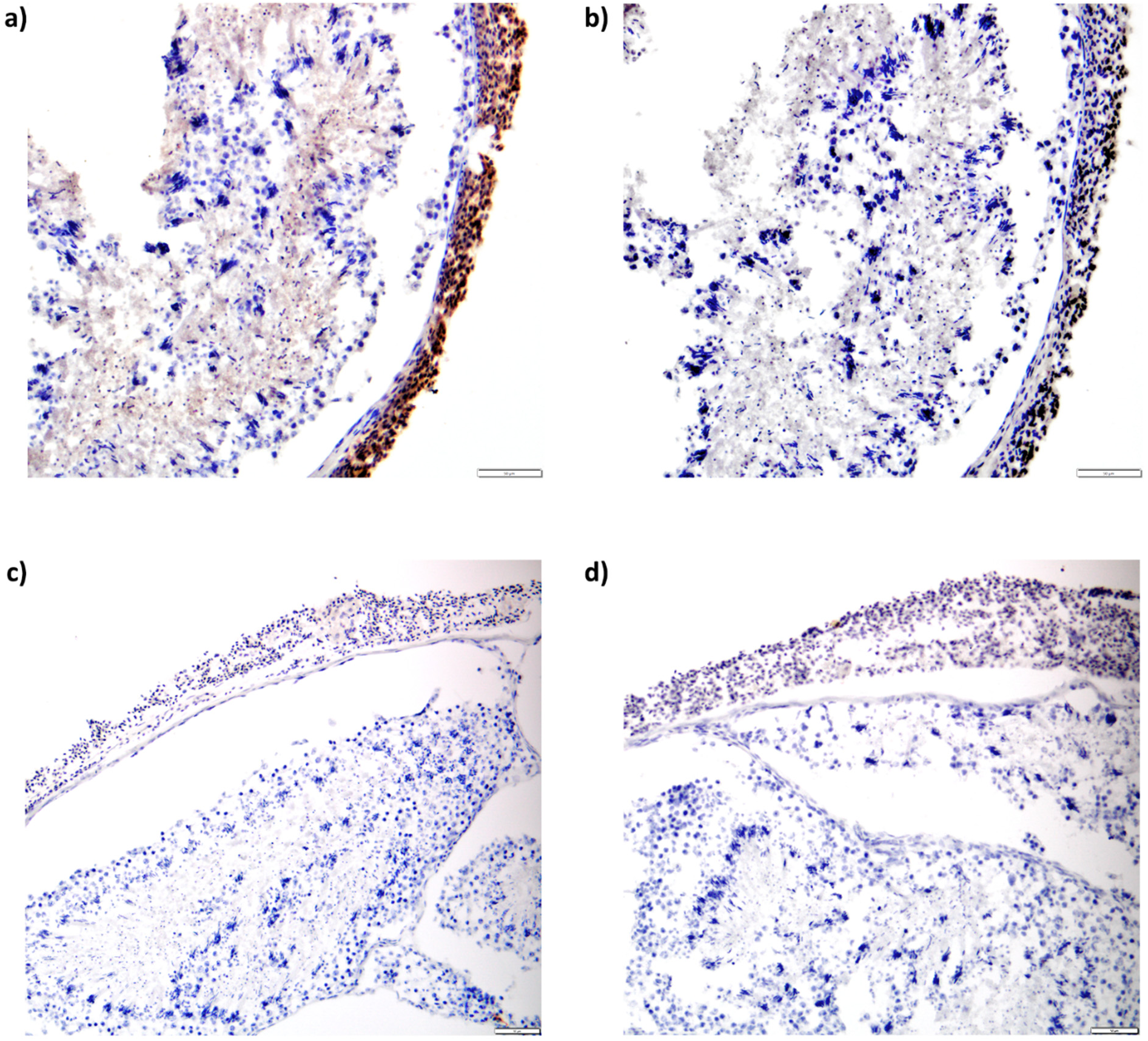
Fgf8 expression in zebra finch testes with positive and negative controls. **a)** Specific expression observed in the tunica albuginea and no expression elsewhere. No expression was observed in the **b)** isotype, **c)** no primary, and **d)** no secondary controls, validating that staining is specific. Scale bar is 50 μm.

**Fig. S7.**
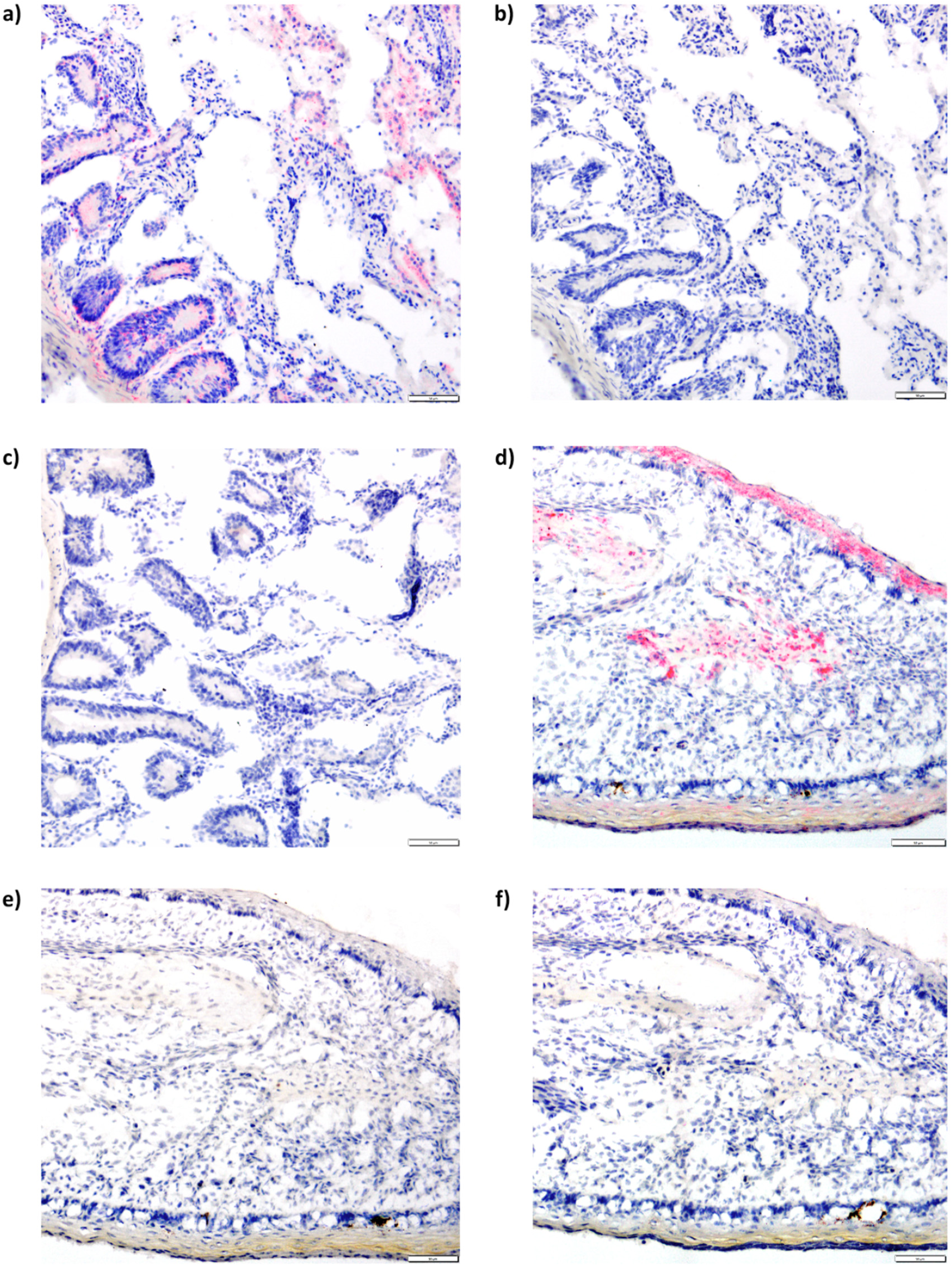
IHH expression in house finch small intestine and lower beak with positive and negative controls. **a)** In small intestine, specific expression observed in areas of mucosa with no expression in muscle. No expression in small intestine tissue was observed in the **b)** isotype, and **c)** no primary controls. **d)** In late stage (HH40) of house finch lower beak, specific expression was observed in chondrocytes and epithelium with no expression in surrounding mesenchyme. No expression in **e)** no primary and **f)** no secondary controls for adjacent beak sections, validating that staining is specific. Scale bar is 50 μm.

**Fig. S8.**
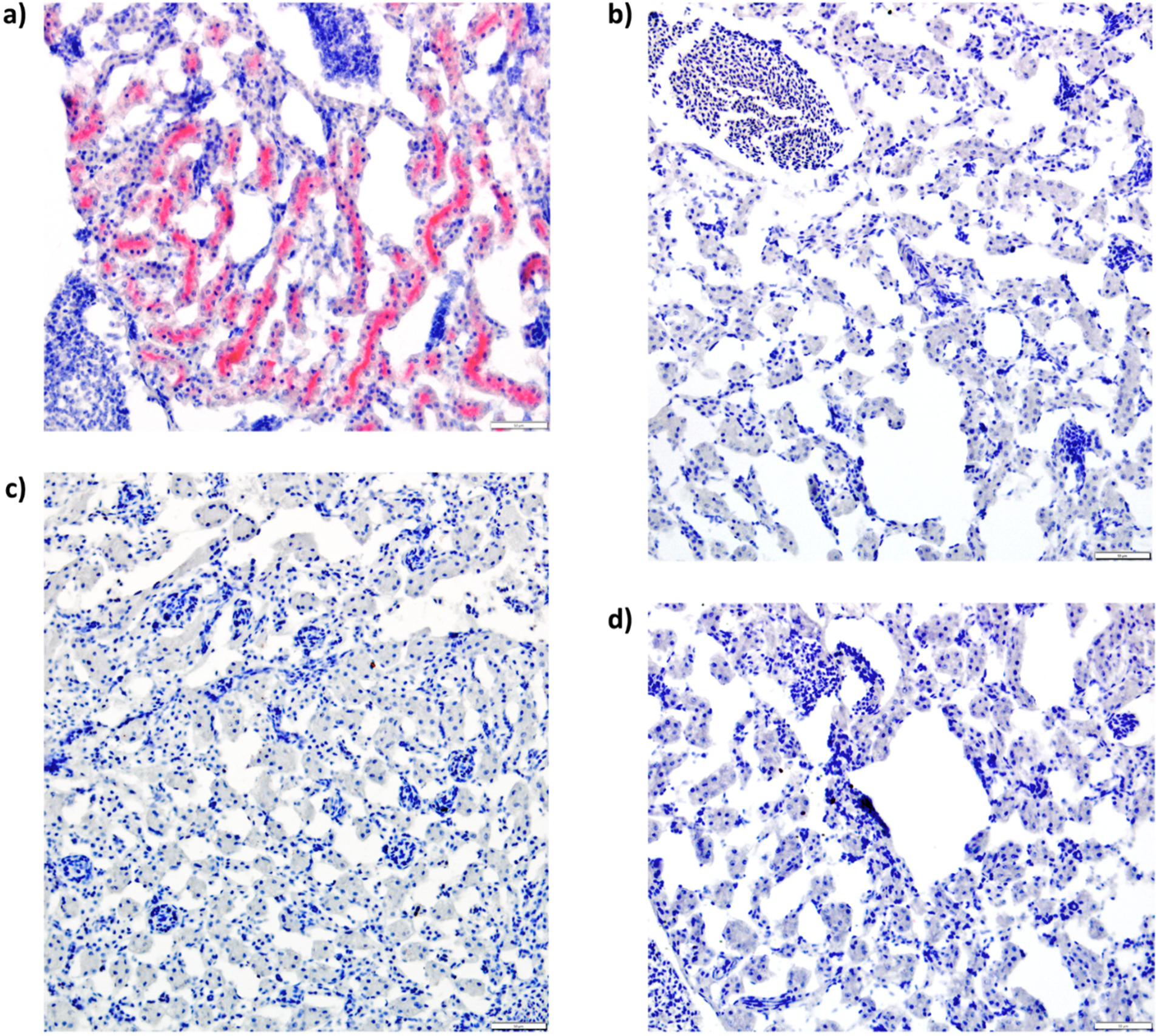
Tgfβ2 expression in house finch kidney with positive and negative controls. **a)** Specific expression observed in extracellular matrix with no expression in glomeruli. No expression was observed in the **b)** isotype, **c)** no primary, and **d)** no secondary controls, validating that staining is specific. Scale bar is 50 μm.

**Fig. S9.**
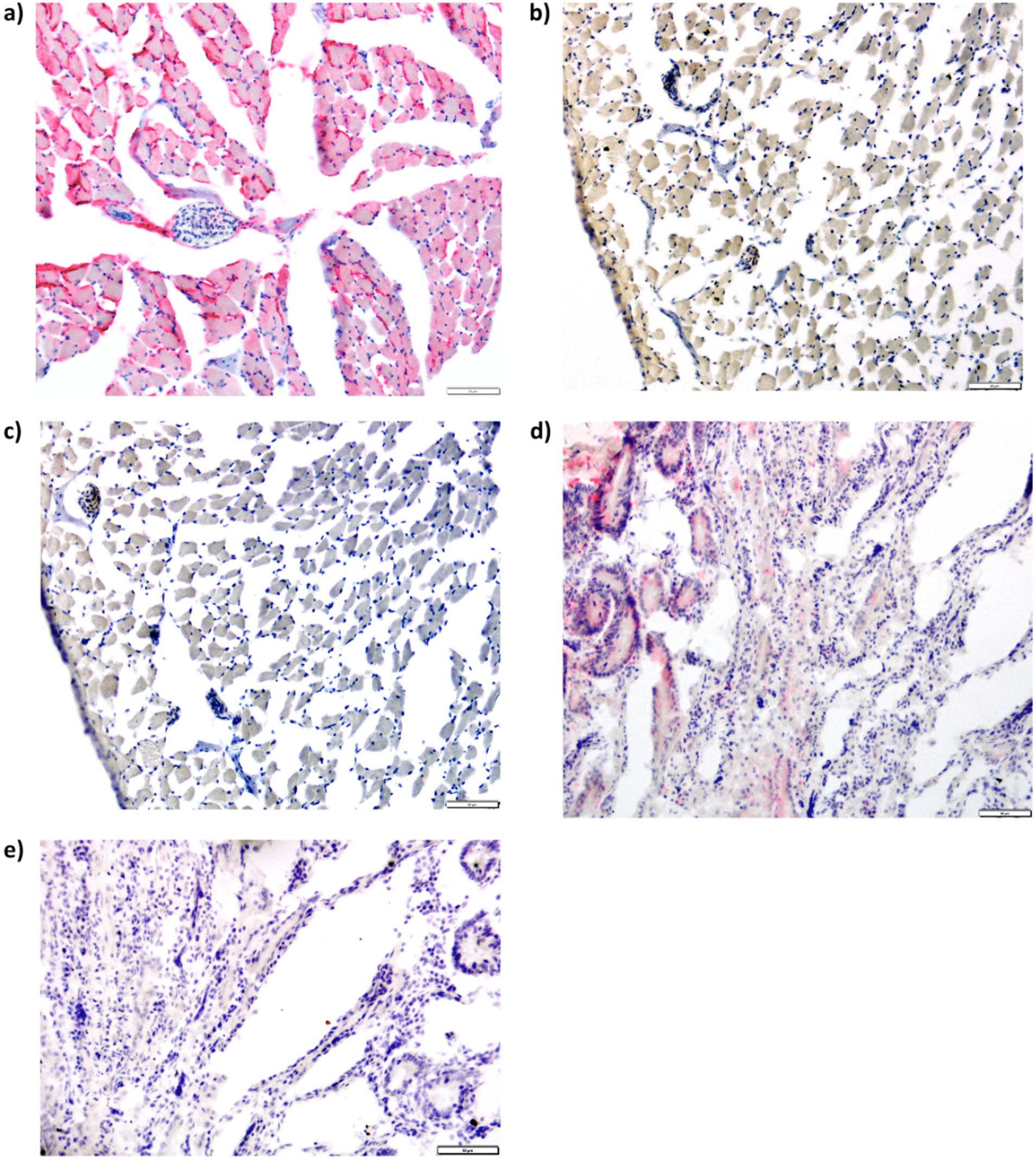
Wnt4 expression in zebra finch thyroid and small intestine with positive and negative controls. **a)** Specific expression observed in follicular epithelium of the thyroid with no expression in colloid and vessels. No expression was observed in the **b)** no primary, and **c)** no secondary controls of thyroid. **d)** In small intestine, specific expression was observed the crypts of Lieberkuhn with no expression in the submucosa. **e)** No expression was observed in the small intestine isotype control, validating that staining is specific. Scale bar is 50 μm.

**Fig. S10.**
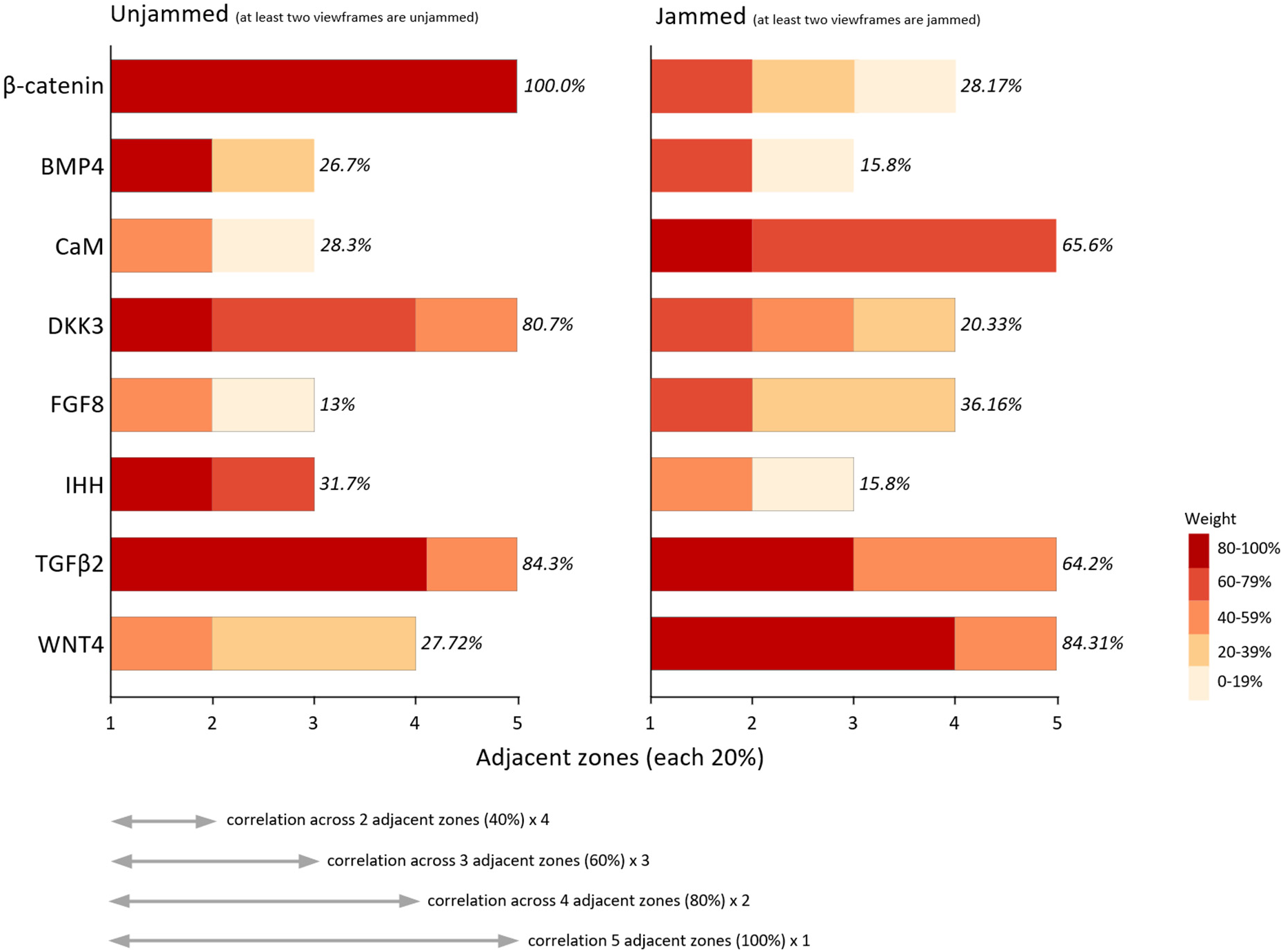
Calculation of spatial reach (correlational span) along anterior-posterior dimension of AOI. Each zone is 20% of the total span. Average span is calculated as a weighed number of adjacent zones with which the expression within a focal zone is significantly correlated (Spearman rank-order correlations). Weight (inset shade scale) is calculated as a number of span distances over which the correlations are possible (e.g., only one span of five zones in length (100%) vs four spans of two zones in length (40%)). Jammed/unjammed state is recorded for a focal (origin) zone only. For example, when β-catenin is expressed in the unjammed tissue within a zone, it is correlated with β-catenin expression across the entire span (all distances are present) of AOI. But when it is expressed in the jammed tissue, its average correlational span is reduced to 28.17%. Zones are considered “unjammed” when at least two viewframes are unjammed and “jammed” when at least two viewframes are jammed.

**Fig. S11.**
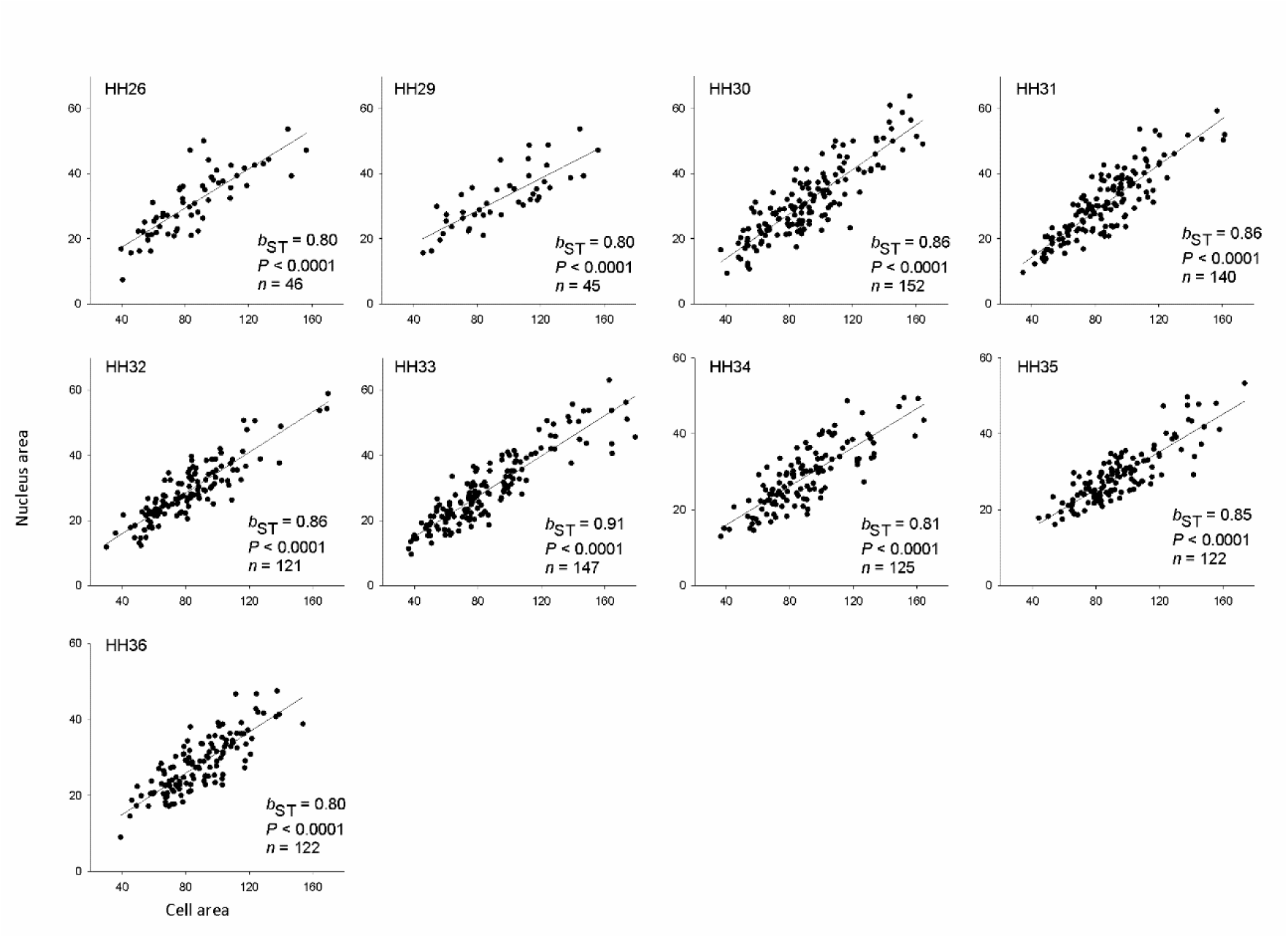
Nucleus area is a suitable proxy for cell area across all developmental stages. Shown are relationships between cell area and nucleus areas (μm^2^) in neural crest mesenchyme across developmental stages as obtained by manual measurements. b_ST_ is standardized regression coefficient.

**Fig. S12.**
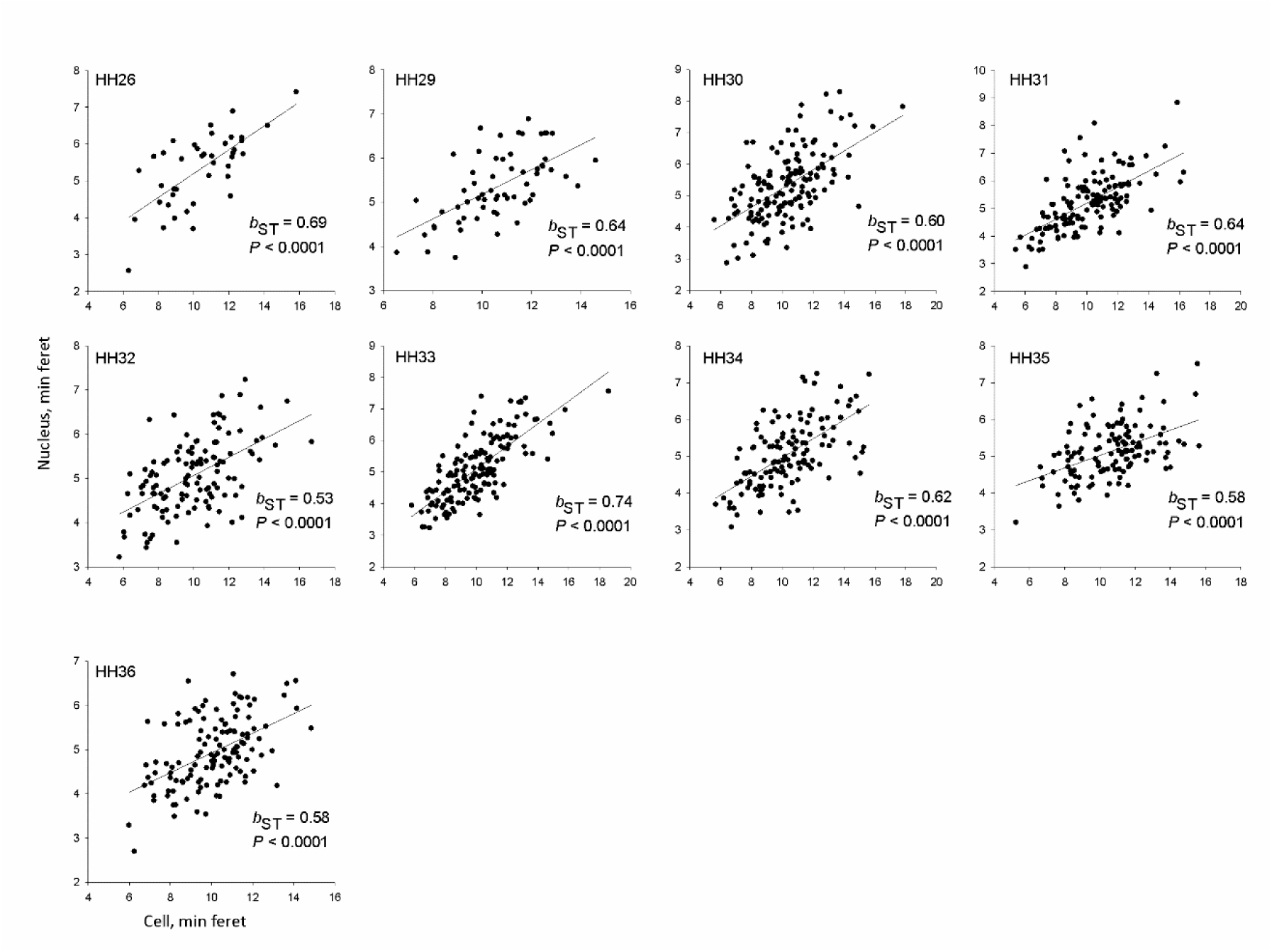
Relationship between cell and nucleus ellipsoid shapes (minimum feret measure shown) in neural crest mesenchyme across developmental stages, manual measurements. b_ST_ is standardized regression coefficient. Sample sizes are in Supplementary Figure S11.

**Supplementary Table S1.**
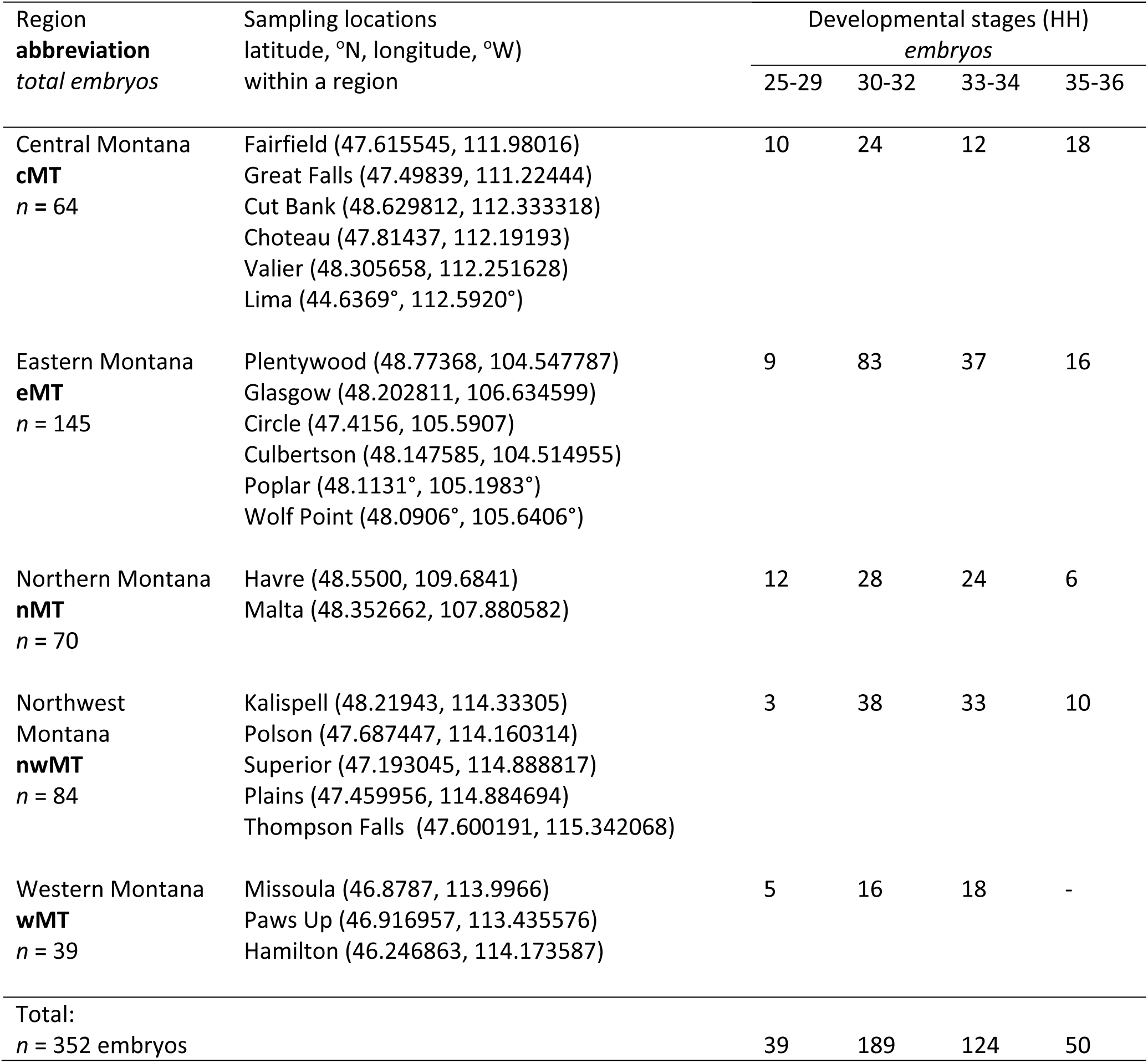
Combined dataset of embryo samples.

| Region<br>**abbreviation**<br>*total embryos* | Sampling locations<br>latitude, °N, longitude, °W)<br>within a region | Developmental stages (HH) |  |  |  |
| --- | --- | --- | --- | --- | --- |
|  |  | *embryos* |  |  |  |
|  |  | 25-29 | 30-32 | 33-34 | 35-36 |
| Central Montana<br>**cMT**<br>*n* = 64 | Fairfield (47.615545, 111.98016)<br>Great Falls (47.49839, 111.22444)<br>Cut Bank (48.629812, 112.333318)<br>Choteau (47.81437, 112.19193)<br>Valier (48.305658, 112.251628)<br>Lima (44.6369°, 112.5920°) | 10 | 24 | 12 | 18 |
| Eastern Montana<br>**eMT**<br>*n* = 145 | Plentywood (48.77368, 104.547787)<br>Glasgow (48.202811, 106.634599)<br>Circle (47.4156, 105.5907)<br>Culbertson (48.147585, 104.514955)<br>Poplar (48.1131°, 105.1983°)<br>Wolf Point (48.0906°, 105.6406°) | 9 | 83 | 37 | 16 |
| Northern Montana<br>**nMT**<br>*n* = 70 | Havre (48.5500, 109.6841)<br>Malta (48.352662, 107.880582) | 12 | 28 | 24 | 6 |
| Northwest Montana<br>**nwMT**<br>*n* = 84 | Kalispell (48.21943, 114.33305)<br>Polson (47.687447, 114.160314)<br>Superior (47.193045, 114.888817)<br>Plains (47.459956, 114.884694)<br>Thompson Falls (47.600191, 115.342068) | 3 | 38 | 33 | 10 |
| Western Montana<br>**wMT**<br>*n* = 39 | Missoula (46.8787, 113.9966)<br>Paws Up (46.916957, 113.435576)<br>Hamilton (46.246863, 114.173587) | 5 | 16 | 18 | - |
| Total:<br>*n* = 352 embryos |  | 39 | 189 | 124 | 50 |

**Supplementary Table S2.** Standardized regression coefficients of AR (mean) vs AR (StdDev) for population groups, developmental stages and zones

| Population | Stage | Zone | $b_{ST}$ | $t$ | $P$ | Population | Stage | Zone | $b_{ST}$ | $t$ | $P$ |
| --- | --- | --- | --- | --- | --- | --- | --- | --- | --- | --- | --- |
| ccMT | 25-31 | 1 | 0.75 | 10.65 | <0.0001 | nwMT | 25-31 | 1 | 0.81 | 10.51 | <0.0001 |
|  |  | 2 | 0.73 | 9.68 | <0.0001 |  |  | 2 | 0.79 | 9.89 | <0.0001 |
|  |  | 3 | 0.77 | 10.06 | <0.0001 |  |  | 3 | 0.71 | 9.55 | <0.0001 |
|  |  | 4 | 0.78 | 11.43 | <0.0001 |  |  | 4 | 0.83 | 10.60 | <0.0001 |
|  |  | 5 | 0.77 | 10.99 | <0.0001 |  |  | 5 | 0.77 | 10.68 | <0.0001 |
|  | 32-33 | 1 | 0.76 | 10.07 | <0.0001 |  | 32-33 | 1 | 0.83 | 11.24 | <0.0001 |
|  |  | 2 | 0.76 | 12.04 | <0.0001 |  |  | 2 | 0.82 | 11.00 | <0.0001 |
|  |  | 3 | 0.81 | 12.54 | <0.0001 |  |  | 3 | 0.76 | 8.71 | <0.0001 |
|  |  | 4 | 0.77 | 10.55 | <0.0001 |  |  | 4 | 0.83 | 12.14 | <0.0001 |
|  |  | 5 | 0.79 | 10.95 | <0.0001 |  |  | 5 | 0.72 | 9.57 | <0.0001 |
|  | 34-36 | 1 | 0.79 | 9.69 | <0.0001 |  | 34-36 | 1 | 0.71 | 10.54 | <0.0001 |
|  |  | 2 | 0.80 | 10.25 | <0.0001 |  |  | 2 | 0.75 | 12.12 | <0.0001 |
|  |  | 3 | 0.85 | 12.53 | <0.0001 |  |  | 3 | 0.71 | 10.19 | <0.0001 |
|  |  | 4 | 0.81 | 10.89 | <0.0001 |  |  | 4 | 0.79 | 14.03 | <0.0001 |
|  |  | 5 | 0.86 | 15.89 | <0.0001 |  |  | 5 | 0.80 | 15.04 | <0.0001 |
| eeMT | 25-31 | 1 | 0.82 | 17.04 | <0.0001 | wwMT | 25-31 | 1 | 0.72 | 8.67 | <0.0001 |
|  |  | 2 | 0.75 | 14.04 | <0.0001 |  |  | 2 | 0.71 | 8.17 | <0.0001 |
|  |  | 3 | 0.70 | 11.57 | <0.0001 |  |  | 3 | 0.71 | 8.20 | <0.0001 |
|  |  | 4 | 0.72 | 12.15 | <0.0001 |  |  | 4 | 0.81 | 8.75 | <0.0001 |
|  |  | 5 | 0.76 | 12.49 | <0.0001 |  |  | 5 | 0.71 | 8.50 | <0.0001 |
|  | 32-33 | 1 | 0.80 | 15.62 | <0.0001 |  | 32-33 | 1 | 0.76 | 11.19 | <0.0001 |
|  |  | 2 | 0.80 | 15.19 | <0.0001 |  |  | 2 | 0.81 | 11.94 | <0.0001 |
|  |  | 3 | 0.84 | 18.05 | <0.0001 |  |  | 3 | 0.72 | 10.67 | <0.0001 |
|  |  | 4 | 0.76 | 12.42 | <0.0001 |  |  | 4 | 0.71 | 10.13 | <0.0001 |
|  |  | 5 | 0.74 | 11.58 | <0.0001 |  |  | 5 | 0.79 | 10.53 | <0.0001 |
|  | 34-36 | 1 | 0.76 | 11.03 | <0.0001 |  | 34-36 | 1 | 0.78 | 9.37 | <0.0001 |
|  |  | 2 | 0.72 | 10.19 | <0.0001 |  |  | 2 | 0.80 | 9.40 | <0.0001 |
|  |  | 3 | 0.72 | 11.19 | <0.0001 |  |  | 3 | 0.72 | 8.92 | <0.0001 |
|  |  | 4 | 0.68 | 10.51 | <0.0001 |  |  | 4 | 0.73 | 9.40 | <0.0001 |
|  |  | 5 | 0.74 | 10.43 | <0.0001 |  |  | 5 | 0.79 | 10.36 | <0.0001 |
| nnMT | 25-31 | 1 | 0.82 | 10.79 | <0.0001 |  |  |  |  |  |  |
|  |  | 2 | 0.82 | 11.89 | <0.0001 |  |  |  |  |  |  |
|  |  | 3 | 0.78 | 8.54 | <0.0001 |  |  |  |  |  |  |
|  |  | 4 | 0.79 | 10.21 | <0.0001 |  |  |  |  |  |  |
|  |  | 5 | 0.74 | 8.38 | <0.0001 |  |  |  |  |  |  |
|  | 32-33 | 1 | 0.72 | 9.13 | <0.0001 |  |  |  |  |  |  |
|  |  | 2 | 0.75 | 10.95 | <0.0001 |  |  |  |  |  |  |
|  |  | 3 | 0.77 | 10.21 | <0.0001 |  |  |  |  |  |  |
|  |  | 4 | 0.75 | 9.85 | <0.0001 |  |  |  |  |  |  |
|  |  | 5 | 0.79 | 10.50 | <0.0001 |  |  |  |  |  |  |
|  | 34-36 | 1 | 0.81 | 10.08 | <0.0001 |  |  |  |  |  |  |
|  |  | 2 | 0.81 | 12.09 | <0.0001 |  |  |  |  |  |  |
|  |  | 3 | 0.84 | 11.28 | <0.0001 |  |  |  |  |  |  |
|  |  | 4 | 0.79 | 10.56 | <0.0001 |  |  |  |  |  |  |
|  |  | 5 | 0.84 | 10.18 | <0.0001 |  |  |  |  |  |  |

**Supplementary Table S3.**
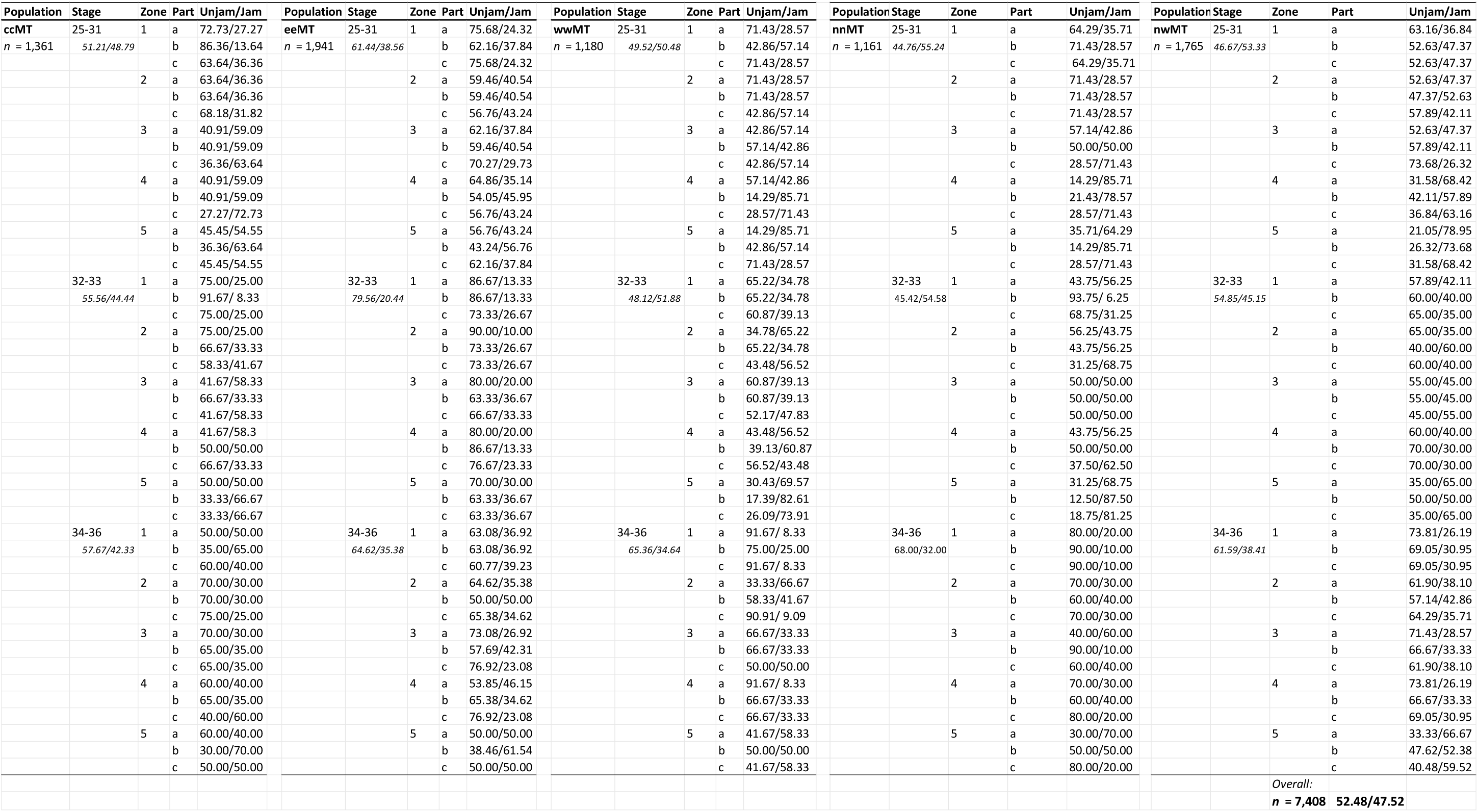
Frequency distribution of unjammed vs jammed cell groups across population groups, developmental stage groups (Stage), zones (1-5) and viewframes (Part). Average ratios are shown for each stage.

**Supplementary Table S4.**
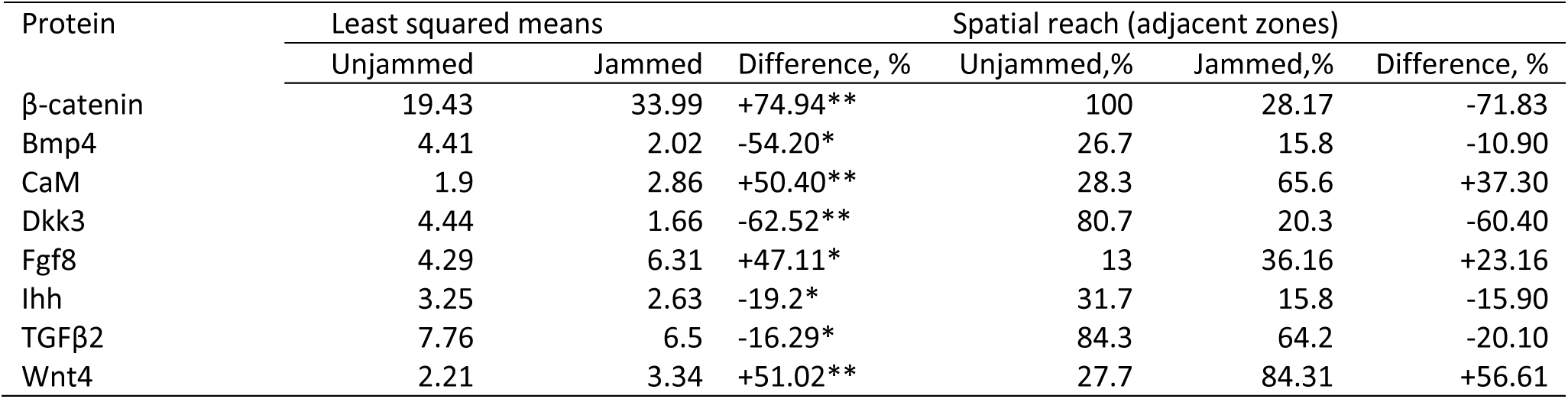
Least squared means (from mixed effects model) for jammed and unjammed cell states, relative difference between these least squared means ((jammed-unjammed)/unjammed)x100%, and protein spatial reach in unjammed and jammed tissues (Fig. S10).

| Protein | Least squared means |  |  | Spatial reach (adjacent zones) |  |  |
| --- | --- | --- | --- | --- | --- | --- |
|  | Unjammed | Jammed | Difference, % | Unjammed,% | Jammed,% | Difference, % |
| β-catenin | 19.43 | 33.99 | +74.94** | 100 | 28.17 | -71.83 |
| Bmp4 | 4.41 | 2.02 | -54.20* | 26.7 | 15.8 | -10.90 |
| CaM | 1.9 | 2.86 | +50.40** | 28.3 | 65.6 | +37.30 |
| Dkk3 | 4.44 | 1.66 | -62.52** | 80.7 | 20.3 | -60.40 |
| Fgf8 | 4.29 | 6.31 | +47.11* | 13 | 36.16 | +23.16 |
| Ihh | 3.25 | 2.63 | -19.2* | 31.7 | 15.8 | -15.90 |
| TGFβ2 | 7.76 | 6.5 | -16.29* | 84.3 | 64.2 | -20.10 |
| Wnt4 | 2.21 | 3.34 | +51.02** | 27.7 | 84.31 | +56.61 |

